# CD74 regulates antitumor immunity in melanoma by reprogramming dendritic cell immunogenicity and migration

**DOI:** 10.64898/2025.12.01.691518

**Authors:** Eleftheria Maranou, Gayoung Park, Pauline Weinzettl, Jonna Alanko, Otto Pulkkinen, Sarah E. Coupland, Marko Salmi, Carlos Rogerio Figueiredo

**Author notes:** Corresponding author. (C.R.F.).

## Abstract

Metastatic cutaneous melanoma remains a major therapeutic challenge, as many patients fail to respond or relapse following immune checkpoint therapy. The function of CD74, originally described as an MHC-II chaperone and receptor for macrophage migration inhibitory factor, remains poorly understood in tumor immunity. In this study, we show that systemic CD74 deletion in a syngeneic melanoma model significantly delayed tumor growth and reshaped the tumor immune microenvironment, and synergized with immune checkpoint blockade. Tumors in *Cd74-/-* mice had more effector and memory CD8+ T cells producing IFN-γ and granzyme B than in control mice. Loss of CD74 promoted expansion of migratory cDC1s with elevated MHC-I and CCR7 expression. *Cd74-/-* professional antigen presenting cells increased cross-priming capacity for tumor-derived antigens in vivo and in vitro. Absence of CD74 induced the expression of CCR7, a key lymph node homing molecule, on cDCs and enhanced their ability to migrate towards the CCL19 ligand in single cell tracking analyses. These findings identify CD74 as a regulator of dendritic cell function and a promising therapeutic target to improve melanoma immunogenicity.

**One Sentence Summary:** CD74 restrains cross-priming and migration capacity of dendritic cells, leading to impaired CD8+ T-cell infiltration and activation in tumor immunity.

## INTRODUCTION

Metastatic cutaneous melanoma is among the most immunogenic malignancies, owing to its elevated tumor mutational burden (TMB) and abundant neoantigen repertoire, features that underpin its sensitivity to immune checkpoint blockade (ICB) therapies (1). Nonetheless, a substantial proportion of patients either fail to respond or relapse following ICB, including dual blockade of CTLA-4 and PD-1 (2). This paradox between immunogenic potential and therapeutic outcome has sparked efforts to elucidate the cellular and molecular determinants within the tumor microenvironment (TME) that shape antitumor immunity and modulate response to therapy.

The TME comprises malignant cells, stromal components, and a heterogeneous immune infiltrate, including dendritic cells (DCs), cytotoxic and regulatory T cells (Tregs), and tumor-associated macrophages (3). Effective tumor immunosurveillance depends on coordinated interactions among these populations (4). For instance, recent work has shown that bidirectional communication between DCs and effector T cells is essential for durable responses to PD-1 blockade (5). Conversely, dominance of immunosuppressive populations such as Tregs correlates with poor outcomes (6), underscoring the need to define molecular regulators that govern immune cell function in metastatic melanoma.

CD74 is a type II transmembrane protein originally characterized as a chaperone for MHC class II (MHC-II) and found to be involved in antigen processing and presentation (7,8). However, beyond its canonical role, CD74 has emerged as a multifunctional regulator of immune activity. Acting as a high-affinity receptor for the macrophage migration inhibitory factor (MIF), CD74 integrates antigen processing with signal transduction pathways that influence inflammation, cell migration, and survival (9–11). Targeting the MIF–CD74 axis has been shown to enhance tumor immunogenicity and restrain melanoma progression in preclinical models (12). Moreover, pharmacological inhibition of MIF was recently reported to synergize with ICB (13), although this effect was only speculated to involve the MIF/CD74 axis, and CD74 involvement was not demonstrated; therefore, the inhibited MIF signals may also arise from its interaction with other receptors (14), highlighting a key limitation of that study.

Despite these findings, the immunological role of CD74 itself, independently of its interaction with MIF, remains poorly understood in tumors, including melanoma. CD74 is expressed across multiple immune cell subsets with distinct functional implications (15). In B cells, the intracellular domain of CD74 acts as a transcriptional regulator, controlling survival and maturation (16). In DCs, CD74 has been reported to participate in MIF family members-mediated signaling, antigen presentation, and has also been found to regulate their migratory capacity (12,17). In Tregs, CD74 regulates cytoskeletal organization, organelle positioning, activation, and Foxp3 expression, thereby promoting their intratumoral accumulation and suppressive functions (18). Recently, CD74 has emerged as a potential biomarker in multiple cancer types, with expression levels variably associated with both favorable and poor outcomes following ICB, including associations with immune infiltration (19,20). Thus far, it remains unclear whether CD74 mainly drives inflammatory or anti-inflammatory immune responses.

Because systemic therapies inhibit their target across the entire organism, constitutive knockout mouse models have been broadly used to define the biological and therapeutic relevance of the target depletion on anti-tumor immunity and tumor progression in vivo. Here, we investigated the contribution of CD74 to melanoma immunity using a *Cd74* knockout (*Cd74-/-*) mouse model. We discovered that CD74 deficiency impaired melanoma progression and reprogrammed the immune architecture of the TME. Tumors in *Cd74-/-* mice exhibited increased infiltration of CD8 T cells, and were characterized by elevated numbers of activated and memory CD8 T cells, alongside heightened number of IFN-γ+ and granzyme B+ CD8 T cells compared to wild-type (*Cd74+/+*) control mice. These changes were accompanied by increased expression of CXCL10 and IFN-α in the TME. In melanoma-bearing *Cd74-/-* mice, DCs displayed elevated surface MHC-I levels, enhanced activation status, and improved cross-priming capacity. CD74 deficiency also led to increased CCR7 expression in DCs and enhanced their migratory capabilities. Collectively, our findings identify CD74 as a critical negative modulator of melanoma immunogenicity and implicate that its loss unleashes a coordinated antitumor immune response.

## RESULTS

### CD74 deficiency remodels the immune architecture of naïve lymph nodes

CD74 mediates tumor-induced immune suppression through MIF interactions in melanoma (12), and breast cancer (21), primarily by limiting DC immunogenicity. Although these findings position CD74 as a potential regulator of myeloid functions, it remains critical for potential therapeutic targeting to understand, whether CD74 also regulates dendritic cell functions in homeostasis and independently from MIF. To decipher the impact of CD74 deficiency in myeloid cells and lymphoid architecture under homeostasis, we used a constitutive *Cd74-/-* mouse model (22).

First, we asked whether complete absence of CD74 influences DC and T cell populations in naïve lymph node, a crucial site of DC and T cell interaction in tumor immunity. Total naïve lymph node cellularity, including immune and stromal cells, was reduced in *Cd74-/-* mice compared to *Cd74+/+* controls (fig. S1A, fig. S2A). Among DC subsets, the frequency of migratory conventional type 1 DCs (cDC1s) was increased in naïve inguinal LNs in *Cd74-/-* mice, while their absolute numbers, as well as those of total and lymphoid-tissue resident cDC1s, remained comparable between genotypes (fig. S1A, fig. S2B). In contrast, the frequencies of cDC2s, and plasmacytoid DCs (pDCs) remained unchanged, and the absolute numbers of cDC2 were reduced in *Cd74-/-* mice (fig. S2B). In *Cd74-/-* mice, CD40 expression was increased in lymphoid-tissue resident cDC1s, CD80 was reduced in migratory cDC1s, and PD-L1 was unaltered across all DC subsets (fig. S2C). Notably, migratory cDC1 showed an upregulation of cell-surface MHC-I, suggesting potentially enhanced antigen presentation capacity (fig. S2D). A trend of MHC-I upregulation was observed in lymphoid-tissue resident cDC1 and cDC2 subsets (fig. S2D). Thus, total DC numbers were unchanged in the naïve lymph node of *Cd74-/-* mice, and the observed differences reflected shifts in subset proportions and increased MHC-I expression per cDC1 rather than an expansion of MHC-I+ cDC1s.

Consistent with the increased frequency of migratory cDC1with higher MHC-I expression, *Cd74-/-* mice exhibited remodeling of their T cell populations. The proportion of CD8+ T cells was increased in *Cd74-/-* mice, whereas the absolute CD8+ T cell numbers remained unchanged (fig. S2E, F). We also observed an expected reduction in CD4+ T cells in the *Cd74-/-* mice, both in frequency and absolute numbers, consistent with the impaired MHC-II-dependent homeostatic maintenance (22,23) (fig. S2E, F). Under steady-state conditions, T cell activation and exhaustion status as well as the proportions of central memory, effector, and naïve T cells were similar in CD4+ and CD8+ T cells in lymph nodes of *Cd74-/-* mice when compared to controls (fig. S2 G, H).

Altogether, these findings suggest that constitutive loss of CD74 remodels the immune architecture of naïve lymph nodes, characterized by reduction in total cellularity, enhanced CD8+ T cell frequency, and reduced CD4+ T cells. Despite the preserved overall DC numbers, *Cd74-/-* mice display higher basal MHC-I, potentially supporting improved immune readiness linked to the increased proportion of immunogenic cDC1 subsets.

### CD74 deficiency controls melanoma growth and enhances accumulation of cytotoxic T cells in the tumor

Building on these findings, we next tested whether the immune alterations observed in naïve lymph nodes of *Cd74-/-* mice enhance anti-tumor immunity in vivo. For that purpose, *Cd74-/-* and *Cd74+/+* control mice were subcutaneously challenged with the ovalbumin-expressing B16 (B16-OVA) melanoma cells (Fig. 1A). *Cd74-/-* mice showed reduced tumor growth compared to *Cd74+/+* controls (Fig. 1B). To determine whether this was associated with changes in the immune landscape at the 20-day endpoint of the experiments, we profiled tumor-infiltrating leukocytes using spectral flow cytometry. This analysis revealed a trend of elevated number of tumor-infiltrating leukocytes, particularly when the leukocyte numbers were normalized per 100 mg of tumor tissue (Fig. 1C, D). The normalization per 100 mg of tumor tissue is used to indicate immune cell density in the TME, allowing comparison of infiltration independently of tumor mass (24). When further analyzing the tumor-infiltrating leukocyte population, we found that CD74 deficiency increased the frequency of CD8+ T cells (Fig. 1E). Analyses of the total numbers of T cells in the tumor (per 100 mg of tumor tissue) confirmed the increased CD8+ T cell infiltration in the *Cd74-/-* mice (Fig. 1F). Although the Tregs numbers also showed a trend of increase in the *Cd74-/-* mice, it was notable that the CD8+ T cells to Treg ratio was increased (Fig. 1G), suggesting a remodeling in the immune landscape that may be associated with enhanced immunosurveillance.

**Figure 1.**
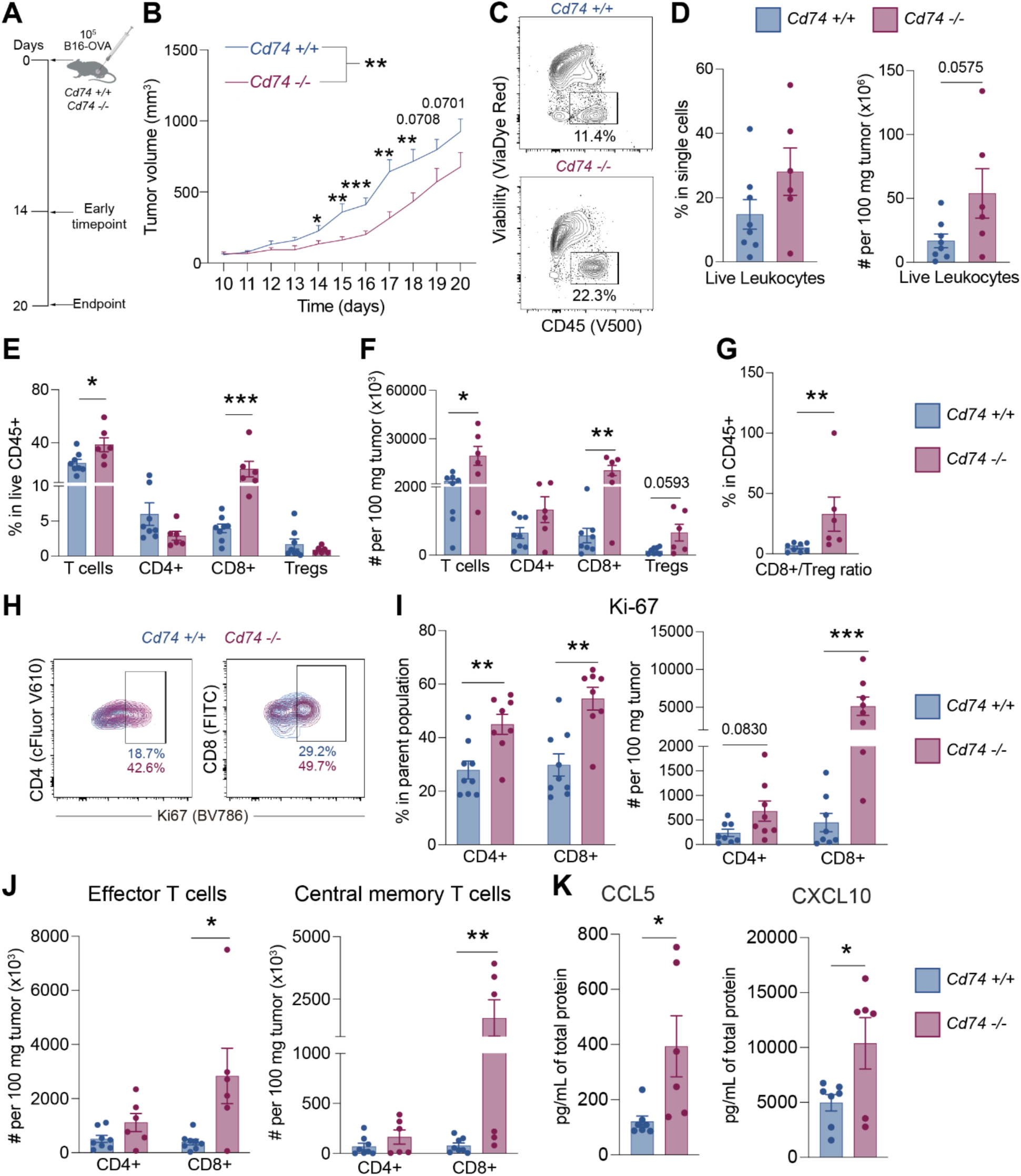
CD74 deficiency attenuates melanoma growth and enhances intratumoral T cell infiltration in syngeneic mice. **(A)** Experimental design for the B16-OVA subcutaneous challenge of 6–10-week-old male *Cd74-/-* and *Cd74+/+* mice. (**B**) B16-OVA tumor growth curves (n=22-31 mice/group from four independent experiments, means + SEM, genotype-factor p-value was determined by two-way ANOVA and p-values for individual time points by multiple unpaired t-test with Welch’s correction). (**C**) Representative flow cytometry contour plots for the frequency of live leukocytes (CD45+) in the TME. (**D**) Frequency and total number of live leukocytes (CD45+) per 100 mg of tumor tissue (n=6-8 mice per group, from two independent experiments, means ± SEM, unpaired t-test). (**E**) Frequencies of all T cells, CD8+ and CD4+ T-cells, and Tregs in the TME (n=6-8 mice per group, from two independent experiments, means ± SEM, Mann-Whitney U test). (**F**) Total number of T cells, CD4+ and CD8+, and Tregs per 100 mg of tumor tissue (n=6-8 mice per group, from two independent experiments, means ± SEM, Mann-Whitney U test). (**G**) Cytotoxic T lymphocyte (CTL, CD8+ T cells) over Tregs ratio calculated from the frequencies of the T-cell populations within live leukocytes (CD45+) (n=6-8 mice per group, from two independent experiments, means ± SEM, Mann-Whitney U test). (**H**) (**I)** Frequency (left) and total number (right) of Ki-67+ cells within the CD4+ and CD8+ T-cell populations per 100 mg of tumor tissue (n=8, from two independent experiments, means ± SEM, Mann-Whitney U test). (**J**) Total number of effector and effector memory (left) and central memory (right) CD4+ and CD8+ T cells per 100 mg of tumor tissue (n=6-8 mice per group, from two independent experiments, means ± SEM, Mann-Whitney U test). (**K**) Intratumoral CCL5 and CXCL10 quantification in B16-OVA tumors in the endpoint of the tumor challenge experiments (n=6-7 mice per group, from two independent experiments, means ± SEM, unpaired t-test). (*\*p≤0.05, **p≤0.01, ***p≤0.001, ****p≤0.0001*)

Moreover, both CD4+ and CD8+ T cells exhibited enhanced frequency of Ki-67+ implying higher proliferation in the TME of the *Cd74-/-* mice (Fig. 1H, I). Notably, an increased numbers of CD8+ T cells infiltrating the TME of *Cd74-/-* mice displayed effector (CD44+ CD62L-) and central memory (CD44+ CD62L+) phenotypes (Fig. 1J, fig. S3A, B) compared to wild-type mice, while no difference was observed in naïve CD4 and CD8 T cells (fig. S3C). Especially, a higher number CD69+ CD8 T cells infiltrated the TME in the absence of CD74 (fig. S3D, E), indicating either enhanced local activation or prolonged tissue residency of these T cells (25,26). When studying the inflammatory status of the TME by intratumoral cytokine quantification, we found a significant increase in the levels of T-cell attracting chemokines CCL5 and CXCL10 in *Cd74-/-* mice (Fig. 1K), supporting the enhanced T cell infiltration observed in these mice.

These findings indicate that CD74 deficiency results in higher proliferation, retention, and memory formation of tumor-infiltrating CD8+ T cells, which associates to attenuated melanoma growth.

### CD74 deficiency promotes intratumoral CD8+ T-cell cytotoxicity and sensitivity to PD-1 blockade

To assess intratumoral T cell functions in the absence of CD74, we analyzed the intracellular expression of IFN-γ and granzyme B (GrB), and the cell-surface expression of PD-1 in the TME-derived lymphocytes in *Cd74-/-* and *Cd74+/+* mice by flow cytometry. Even though the frequencies of IFN-γ+ CD4 and CD8 T cells did not show a significant difference between the genotypes, analyzing the absolute number of cells (per 100 mg of tumor) revealed a significant increase in IFN-γ+ CD8 T cells in *Cd74-/-* mice (fig. S4A and B, Fig. 2A). The number of CD8+ T cells expressing GrB also showed an increase in tumors of *Cd74 -/-* mice, while only frequencies of GrB+ CD4 showed a trend for increase (Fig. 2A, fig. S4C and D). We also observed that the absolute number of PD-1+ CD8 T cells was increased in the TME of *Cd74-/-* mice (Fig. 2B). Together, these findings suggest that absence of CD74 supports increased infiltration or accumulation of activated and PD-1+ cytotoxic CD8 T cells.

**Figure 2.**
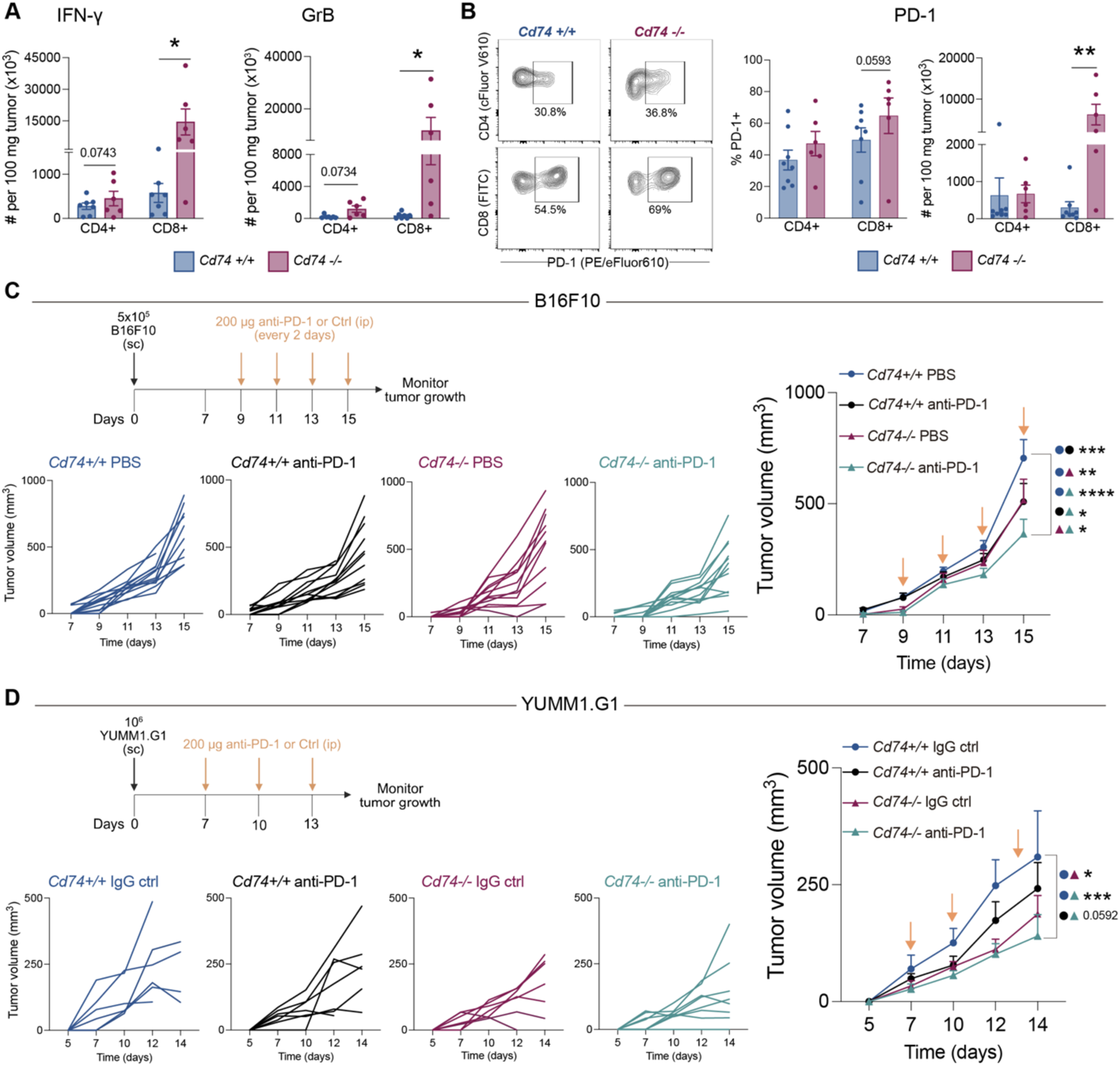
CD74 deficiency leads to increased infiltration by activated CD8+ T cells in their TME and potentiates anti-PD1 therapy. (**A**) Total number (per 100 mg of tumor tissue) of IFN-γ+ (left) and GrB+ (right) CD4+ and CD8+ T cells in the TME (n=6-7 mice per group, from two independent experiments, means ± SEM, unpaired t-test for frequency, Mann-Whitney U test for total cell amounts). (**B**) Representative flow cytometry contour plots for analyzing the frequency (left) and total number (right; per 100 mg of tumor tissue) of PD-1+ CD4+ and CD8+ T cells in the TME (n=6-7 mice per group, from two independent experiments, means ± SEM, unpaired t-test for frequency, Mann-Whitney U test for total cell amounts). (**C-D**) B16F10 (C) and YUMM1.G1 (D) tumor-bearing mice were treated with PBS/isotype (IgG) control or anti-PD1 at the indicated time points (B16F10: days 9, 11, 13, 15; YUMM1.G1: days 7, 10, 13) (n=12 mice/group for B16F10 and n=6-8 mice/group for YUMM1.G1 from two independent experiments, means + SEM, p-values for day 15 (B16F10) and day 14 (YUMM1.G1) were determined by two-way ANOVA with Tukey’s multiple comparisons test). (*\*p≤0.05, **p ≤0.01, ***p≤0.001, ****p≤0.0001*)

Given the elevated number of PD-1+ CD8 T cells within the TME of *Cd74-/-* mice, we investigated whether CD74 deficiency modulates responsiveness to PD-1 blockade. Tumor growth was compared between *Cd74-/-* and *Cd74+/+* mice treated with anti-PD-1 using B16F10 and YUMM1.G1 melanoma models, both of which develop resistance to anti-PD-1 immunotherapy. In the B16F10 model, anti-PD-1 significantly delayed tumor progression in *Cd74-/-* compared to *Cd74+/+* mice (Fig. 2C), indicating enhanced therapeutic efficacy in the absence of CD74. Similarly, in the YUMM1.G1 model, anti-PD-1 resulted in a marked reduction in tumor growth in *Cd74-/-* mice relative to *Cd74+/+* controls (Fig. 2D), despite the intrinsic resistance of this model to PD-1 blockade. Across both melanoma models, the most pronounced tumor control was observed in Cd74-/- mice receiving PD-1 blockade.

Collectively, these results demonstrate that CD74 deficiency synergizes with PD-1 blockade to enhance anti-tumor efficacy, even in melanoma models with limited baseline responsiveness to ICB.

### CD74 deficiency elevates MHC-I on DCs with minimal impact on their activation phenotype

The intratumoral T cell findings led us to hypothesize that CD74 deficiency may alter DCs composition or functions to favor immune activation locally in the TME and tdLN. Even though the frequencies of DCs in TME remained unchanged between the genotypes (Fig. 3A, B), there was a trend for increased number of DCs (per 100 mg of tumor tissue) in *Cd74-/-* mice (Fig. 3A, B). Subset analysis of TME showed no differences in the abundance of cDC subsets between genotypes (Fig. 3C). Expression of CD40 was comparable between *Cd74-/-* and *Cd74+/+* control mice across all DC subsets (Fig. 3D). However, the proportion of migratory cDC1s expressing PD-L1 was higher in the TME of *Cd74-/-* compared to *Cd74+/+* mice (Fig. 3E). In addition, these cells showed a trend toward increased PD-L1 expression, as indicated by the median fluorescence intensity (MFI) (Fig. 3E). We also observed that the frequency of MHC-I+ cDC1s and their migratory subset showed a trend for an increase, and that MHC-I expression level on both cell types was enhanced in *Cd74-/-* mice (Fig. 3F).

**Figure 3.**
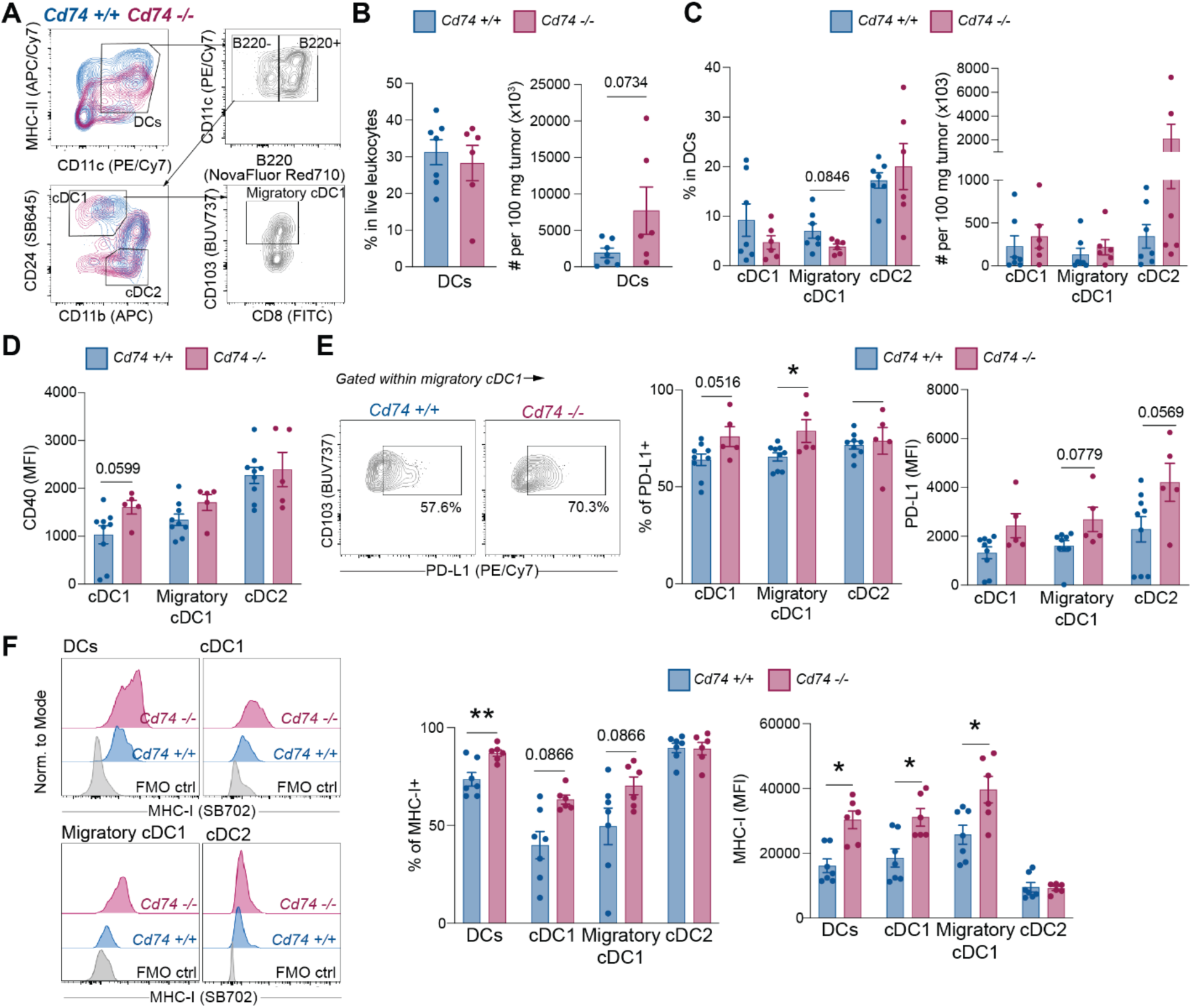
CD74 deficiency elevates MHC-I on cDC1s in the TME. (**A**) Gating strategy used for the discrimination of DCs and their subpopulations. (**B and C**) Frequency and absolute number of (B) DCs and (C) DC subsets per 100 mg of tumor tissue (n=6-7 mice per group, from two independent experiments, means ± SEM, Mann-Whitney U test). (**D**) Median fluorescence intensity (MFI) of CD40 (n=5-9 mice per group, from two independent experiments, means ± SEM, Mann-Whitney U test). (**E**) (From left to right) Representative flow cytometry contour plots for analyzing the frequency of PD-L1 on migratory cDC1, and frequency as well as MFI of PD-L1 in DC subsets in the TME (n=5-9 mice per group, from two independent experiments, means ± SEM, Mann-Whitney U test). (**F**) (Left) Representative flow cytometry histograms indicating the cell surface expression of MHC-I in different DC subsets in the TME. (Right) Frequency and MFI of MHC-I calculated within DC subsets in the TME (n=6-7, from two independent experiments, mean ± SEM, Mann-Whitney U test). (*\*p≤0.05, **p≤0.01, ***p≤0.001, ****p≤0.0001*)

Interestingly, *Cd74-/-* tumor-bearing mice showed a trend for increased pDCs frequency accompanied by an increase in the total number of pDCs (per 100 mg of tumor tissue) (fig. S5A). The co-stimulatory molecule CD80 was expressed on higher proportion of pDCs and at increased level in *Cd74-/-* mice, suggesting enhanced activation of this subset (fig. S5B). Consistently, the concentration of IFN-α, primarily derived from pDCs (27), was elevated in the TME of *Cd74-/-*mice (fig. S5C). The frequency MHC-I+ pDCs and the expression level MHC-I on pDCs in the TME of *Cd74-/-* mice were increased (fig. S5D).

Since type I interferons are known to promote the migration of cDC1s to tumor-draining lymph nodes (tdLNs) and facilitate their transport of tumor-derived antigens (28), we hypothesized that the increased IFN-α in the TME could have promoted earlier or continuous migration of antigen-loaded cDC1s from the tumor to the tdLNs, which together with the enhanced MHC-I expression potentially shape systemic immune responses by supporting antigen presentation in lymphoid organs. To explore this possibility, we next examined the myeloid compartment in the tdLNs in different stages of tumor progression.

In the early timepoint (d14) after tumor challenge, *Cd74-/-* mice displayed increased frequency of total cDC1, migratory cDC1s, and lymphoid-tissue resident cDC1, while absolute numbers of migratory cDC1 where comparable between the genotypes and resident cDC1 showed a trend toward lower absolute numbers (Fig. 4A). Notably, the absolute numbers of cDC2 were reduced in tdLNs of *Cd74-/-* mice, despite their unchanged frequency. The total tdLNs cellularity did not differ between the genotypes (fig. S6A). pDCs did not show difference in their proportions between the genotypes (fig. S5E). By the endpoint (d20), the proportions of different DC subpopulations changed with migratory cDC1s showing a trend for decrease in *Cd74-/-* tdLNs, (Fig. 4B). In contrast the absolute number of cDC1, their subsets, and cDC2 did not differ between *Cd74+/+* and *Cd74-/-* mice. The frequency of pDCs increased in the *Cd74-/-* tdLNs at the endpoint (fig. S5F). The MHC-I levels of cDC1s, specifically their lymphoid-tissue resident subset, were elevated in the tdLNs of *Cd74-/-* mice at the endpoint (Fig. 4C). Therefore, CD74 deficiency transiently leads to reorganization of DCs in tdLNs, with a higher proportion of cDC1 populations in the tdLNs during early tumor growth, rather than expansion of the cDC1 subsets. This potentially reflects altered dendritic cell trafficking in the absence of CD74, which may contribute to the altered immune environment observed in the tumors of *Cd74-/-* mice.

**Figure 4.**
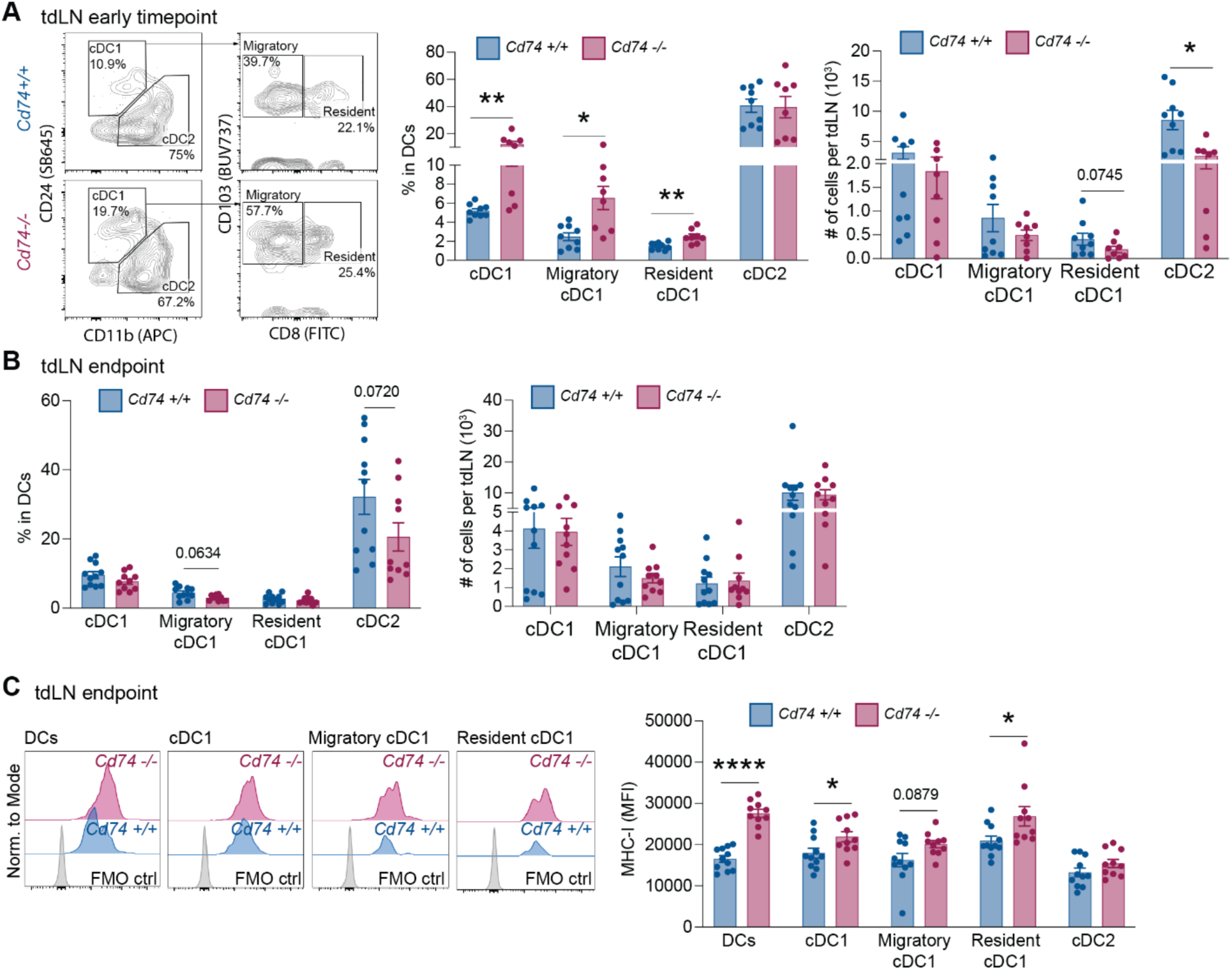
Altered migration kinetics and MHC-I expression of cDC1s in the tdLN of CD74-deficient tumor-bearing mice. (**A and B**) Representative flow cytometry contour plot for analyzing cDC1 subsets (the percentages inside contour plots are within parent populations) (left), frequency and absolute numbers of DC subsets (right) calculated within DCs in tumor draining lymph nodes on (A) day 14 (early timepoint) and (B) day 20 (endpoint) after subcutaneous melanoma cell inoculation (n=8-11 mice per group, from two independent experiments, mean ± SEM, Mann-Whitney U test). (**C**) Representative flow cytometry histograms of MHCI expression (left) and MFI of MHCI (right) in different DC subsets in the tdLN on day 20 (endpoint) of the tumor challenge. (n=8-11, from two independent experiments, mean ± SEM, Mann-Whitney U test). (*\*p≤0.05, **p≤ 0.01, ***p≤0.001, ****p≤0.0001*)

Analysis of T cells in tdLNs at early timepoint and endpoint during melanoma progression revealed distinct differences in CD4 and CD8 T cell dynamics between *Cd74-/-* and *Cd74+/+* mice. In early time point, *Cd74-/-* mice showed a trend for increased T cell frequency in their tdLNs (fig. S6B). Even though this difference was not observed in the absolute numbers of T cells, *Cd74-/-* mice had a 4-fold increase in their T cell absolutes numbers from the early timepoint to the endpoint of the experiment, similar to *Cd74+/+* mice (fig. S6B). At both timepoints, *Cd74+/+* mice had higher frequency and absolute numbers of CD4+ T cells (fig. S6C), whereas in *Cd74-/-*mice the CD8+ T cell population was dominant (fig. S6D). Within-genotype analysis showed that the CD8+ T cell frequency declined from early to endpoint, while the opposite was observed when comparing the absolute numbers of CD8 T cells (fig. S6D). These data suggest that the early and transient reorganization of the DC compartment precedes and might be associated with a sustained expansion of CD8+ T cells in *Cd74-/-* mice. At early timepoint, higher frequency of CD4+ T cells expressed CD69 in *Cd74-/-* tdLNs, with the difference between the genotypes becoming more pronounced by the endpoint (fig. S6B). In contrast, in *Cd74+/+* mice the frequency of CD69+ CD8 T cells did not increase between the early and late timepoints (fig. S6B). In addition, the tdLNs of *Cd74-/-* were enriched with PD-1+ CD4 and CD8 T cells in early timepoint, but only by increased PD-1+ CD4 T cells in endpoint (fig. S6C). The increased presence of PD-1+ T cells in the tdLNs of *Cd74-/-* mice may contribute to the enhanced responsiveness to PD-1 blockade we observed in our anti-PD1 treatment (Fig. 2C and D), supporting functional cooperation between CD74 deficiency and PD-1 blockade.

### CD74 deficiency enhances CD8 T cell priming in vivo

Our phenotyping data at the endpoint of tumor challenge indicate that MHC-I levels on cDC1 subsets are influenced by CD74 loss. The elevated expression of MHC-I on DCs together with the increased intratumoral CD8 T cells could potentially suggest improved antigen cross-presentation performed by the CD74-lacking DCs. Hence, we next examined the role of CD74 in cross-presentation of cancer cell-associated antigens.

We first used an in vivo proliferation assay. We adoptively transferred CellTrace Violet (CTV)-labeled TCR-transgenic T cells (OT-I) recognizing SIINFEKL, the MHC-I restricted peptide derived from ovalbumin (OVA), in *Cd74-/-* and *Cd74+/+* control mice. Five days after the immunization of *Cd74-/-* and *Cd74+/+* controls with OVA-expressing apoptotic B16-melanoma cells (Fig. 5A), OT-I T cells were found in higher frequency in the peripheral blood, tdLNs, and spleens of *Cd74-/-* mice (Fig. 5B, C, F, G, fig. S7A, B, fig. S8A). Notably, the OT-I cells in the peripheral blood and tdLNs had proliferated more in *Cd74-/-* mice, when assessed by analyzing the frequencies of cells expressing a SIINFEKL dextramer-binding T-cell receptor and CTV^low^CD44^high^ OT-I cells, respectively (Fig. 5D-E, 5H-I), while no difference in OT-I cell-proliferation was observed in the spleens (fig. S8B).

**Figure 5.**
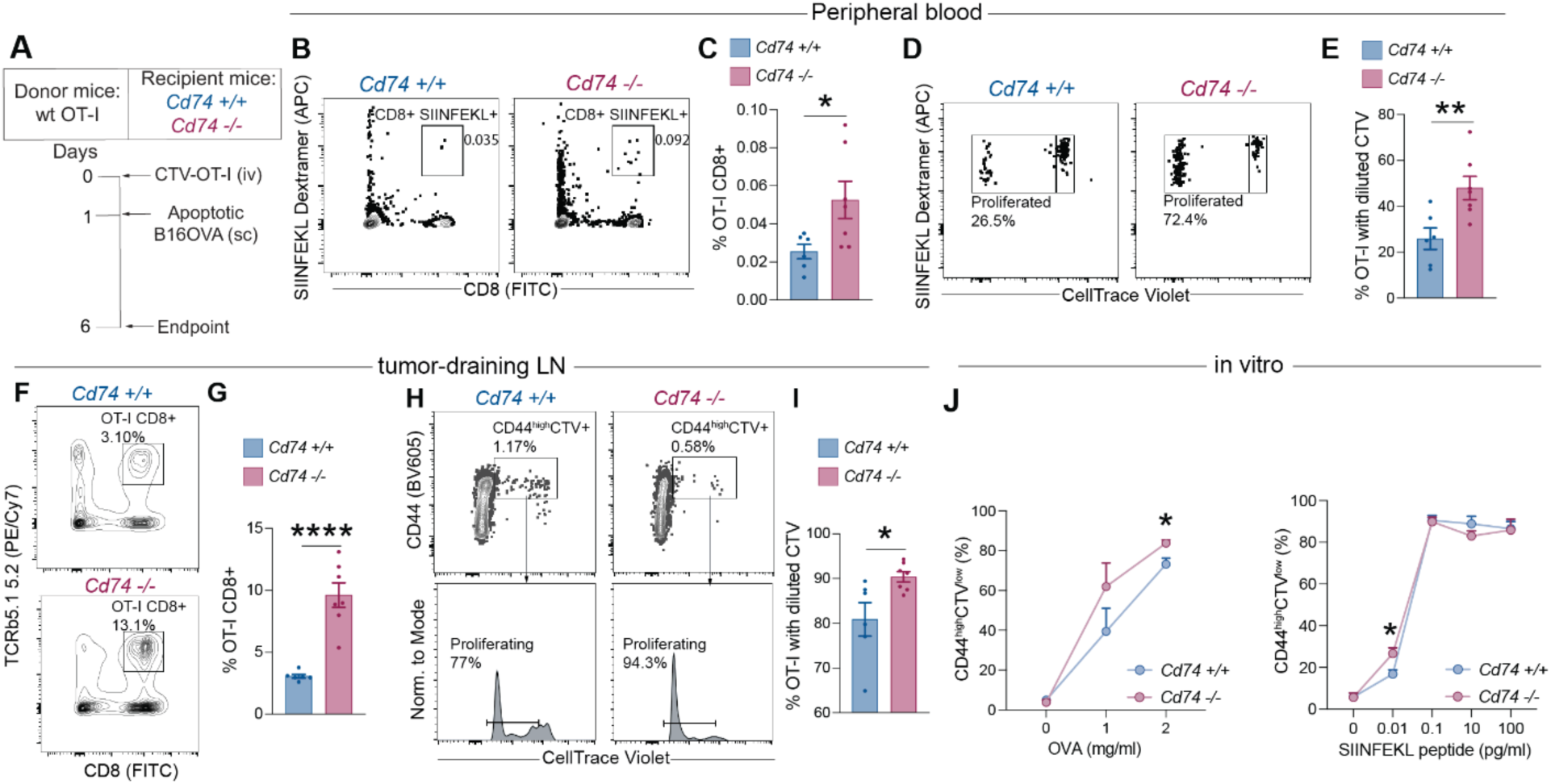
Antigen-presenting cells of *Cd74-/-* mice demonstrate enhanced cross-priming capabilities. (**A**) Experimental design of in vivo cell-associated antigen cross-presentation assay. (**B**) Representative flow cytometry contour plots showing the frequency of OT-I CD8+ T cells (SIINFEKL Dextramer+ CD8+) in the peripheral blood on day 6, gated within single cells. (**C**) Frequency of OT-I CD8+ T cells in the peripheral blood on day 6 (n=6-7 mice per group, from two independent experiments, mean ± SEM, unpaired t-test). (**D**) Representative flow cytometry dot plots indicating the proliferated (diluted CTV) OT-I CD8+ T cells (SIINFEKL Dextramer+ CD8+) in the peripheral blood, gated within OT-I CD8+ T cells. (**E**) Frequency of proliferated OT-I CD8+ T cells in the peripheral blood on day 6 (n=6-7 mice per group, from two independent experiments, mean ± SEM, unpaired t-test). (**F**) Representative flow cytometry contour plots showing the frequency of OT-I CD8+ T cells (TCRb5.1 5.2+ CD8+) in the tdLNs on day 6, gated within live leukocytes. (**G**) Frequency of OT-I CD8+ T cells in the tdLNs on day 6 (n=6-7 mice per group, from two independent experiments, mean ± SEM, unpaired t-test). (**H**) Gating strategy used for the discrimination of activated OT-I CD8+ T cells expressing CD44, positive for CTV, and subsequent gating of activated OT-I CD8+ T cells with diluted CTV (proliferating cells) in tdLNs. (**I**) Frequency of activated OT-I CD8+ T cells with diluted CTV in the tdLNs on day 6 (n=6-7 mice per group, from two independent experiments, mean ± SEM, unpaired t-test). (**J**) (Top) Frequency of CD44^high^CTV^low^ OT-I CD8+ T cells after a 3-day incubation with ovalbumin-pulsed *Cd74-/-* and *Cd74+/+* FL-DCs (n=4 independent experiments, mean ± SEM, unpaired t-test). (Bottom) Frequency of CD44^high^CTV^low^ OT-I CD8+ T cells after a 3-day incubation with SIINFEKL-pulsed *Cd74-/-* and *Cd74+/+* FL-DCs (n=3-5 independent experiments, mean ± SEM, unpaired t-test). (*\*p≤0.05, **p≤0.01, ***p≤0.001, ****p≤0.0001*)

The in vivo findings suggested an enhanced cross-priming capacity of DCs in tdLN of *Cd74-/-*mice. To exclude the contribution from non-DC cell types, we performed an in vitro antigen presentation assay using FLT3L-generated bone marrow-derived DCs (FL-DCs) from *Cd74-/-* and *Cd74+/+* mice. The FL-DCs were pulsed with different concentrations of either chicken OVA, which requires intracellular antigen processing or with the SIINFEKL peptide, which directly binds to cell-surface MHC-I. After antigen pulsing, the FL-DCs were co-cultured with CTV-labeled OT-I T cells, and T cell proliferation was assessed after 3 days. OVA-pulsed (2 mg/mL) *Cd74-/-* FL-DCs induced enhanced OT-I proliferation compared to *Cd74+/+* FL-DCs (Fig. 6J, fig. S7C). Similar result was observed when FL-DCs were pulsed with the lowest concentration of SIINFEKL (0.01 pg/mL) (Fig. 6J).

**Figure 6.**
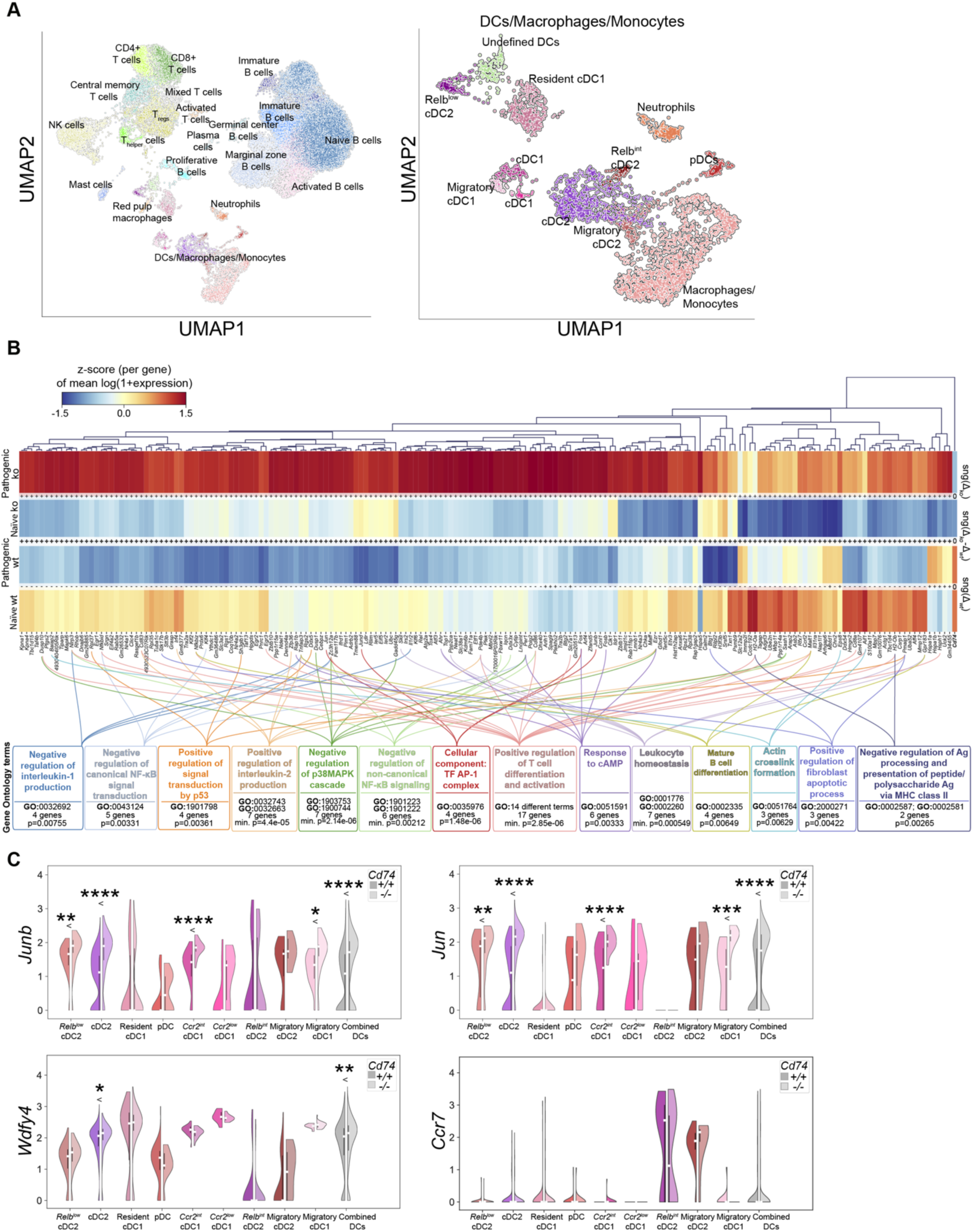
scRNA-seq analysis revealed an association between Cd74 and members of the AP-1 complex. (A) UMAPs showing the cell populations identified in the spleens of naïve and tumor-bearing Cd74-/- and Cd74+/+ mice**. (B)** Heat map representation of the DEGs list using genewise Z-score normalization derived from DC pseudobulk interaction DEA and the immune relevant GO terms related to the DEGs. Naïve wt and ko refers to naïve Cd74+/+ and Cd74-/- mice, while pathogenic wt and ko refers to B16-OVA-bearing Cd74+/+ and Cd74-/- mice, respectively. Because of the interaction effects statistical model, DEGs have resulted from the highest differential expression between Cd74+/+ and Cd74-/- mice in response to B16-OVA (pathogenicity). The plus (+) and minus (-) symbols between naïve wt (Cd74+/+) and pathogenic wt (B16-OVA-bearing Cd74+/+) as well as between naïve ko (Cd74-/-) and pathogenic ko (B16-OVA-bearing Cd74-/-) show the direction of expression change in Cd74+/+ and Cd74-/-, sng(Δwt) and sng(Δko), respectively. The bold plus (+) symbols in the middle of the heatmap indicate that the total change sng(Δko- Δwt) was increased differential expression in the Cd74-/- as compared to Cd74+/+ when pathogenicity (B16-OVA tumors) is introduced. Below the heatmap, the colorful boxes are indicating the GO terms with at least 5% of the pathway genes represented in DEGs derived from the interaction effects DEA in the spleen samples of the naïve and B16-OVA-bearing mcie (p_corrected<0.01, FDR: Benjamini-Hochberg). (**C**) Expression levels of Jun, Junb, Wdfy4, and Ccr7 in all DC subsets identified in the tdLN of Cd74-/- and Cd74+/+ mice (Mann–Whitney U test with Bonferroni correction). (*\*p≤0.05, **p≤0.01, ***p≤0.001, ****p≤0.0001*)

To further exclude potential genotype-specific differences in antigen uptake and proteolysis, we utilized the DQ-OVA assay, which measures fluorescence signal derived from quenched DQ-OVA after proteolytic degradation of the internalized antigen. We observed that *Cd74-/-* resting or LPS-matured FL-DCs exhibited similar DQ-OVA processing compared to their wild-type counterparts (fig. S9A, B). Thus, these data suggest that lack of CD74 in DCs leads to enhanced cross-presentation of neoantigens without affecting the ability of DCs to proteolytically process internalized antigens.

### CD74 deficiency is associated with upregulation of *c-Jun/JunB* at transcriptome level in response to melanoma

Building on these findings, we performed single-cell RNA sequencing (scRNA-seq) to explore the molecular pathways underlying the observed alterations in DC functions in *Cd74-/-* mice. We analyzed lymph nodes and spleens of *Cd74-/-* as well as *Cd74+/+* naïve and B16-OVA-bearing mice. Tumor samples were amenable to analyses due to low immune cell number in the *Cd74+/+* sample and the differential expression analysis (DEA) was not performed between the naïve and tumor-draining lymph node samples due to the low number of DCs. We first annotated cell clusters and identified DC populations in the spleen and lymph node samples using subset-specific signature genes (29–32) (Fig. 6A, fig. S10A-C). The aggregated DC clusters of the spleen samples, excluding macrophages and monocytes, were subjected to DEA between naïve and tumor-bearing *Cd74+/+* versus naïve and tumor-bearing *Cd74-/-* mice, revealing differentially expressed genes (DEGs) from the interaction of the two genotypes in steady-state (naïve samples) and cancer (pathogenic samples) (Fig. 6B). Therefore, we used interaction effects statistical model in order to clarify if the genotype affects the response to disease (33).

To better understand the biological processes and pathways underlying the transcriptional changes observed in the absence of CD74, we performed gene ontology (GO) enrichment analysis on the DEGs identified from the interaction DEA. This analysis highlighted several immune-related pathways including negative regulation of interleukin-1 production and NF-κB signaling (Fig. 6B). Moreover, the AP-1 transcription factor complex was enriched as cellular component based on differential expression of *Jun*, *Junb*, *Fos*, and *Jund* (Fig. 6B). Notably, both *Jun* and *Junb* showed significant upregulation across cDC2, cDC1, migratory cDC1 and total DC populations in the spleen of *Cd74-/-* B16-OVA-bearing mice (Fig. 6C). *Jun* and *JunB* have been identified as essential transcription factors for lymphoid-tissue resident cDC1 function and maintenance in the spleen (34), and to be associated to *Wdfy4* expression, essential for cell-associated antigen cross-presentation by cDC1s and cDC2s (34,35). We observed an upregulation of *Wdfy4* only when combining all DC clusters together and in cDC2 cluster, with a trend for increased expression in migratory cDC1s in the spleen of *Cd74-/-* B16-OVA-bearing mice (Fig. 6C). Moreover, *Asb2*, *Gpr183*, and *Eps8* were among the DEGs and assigned to GO: 0036336, dendritic cell migration (fig. S10D; GO term p=0.0125). Even though dendritic cell migration was highlighted from the GOA, *Ccr7* mRNA expression in the spleen did not show significant differences between *Cd74-/-*and *Cd74+/+* B16-OVA-bearing mice (Fig. 6C). In the tdLN samples, *Junb* was upregulated in cDC2 cells of *Cd74+/+* mice and *Jun* was upregulated in combined DCs of *Cd74-/-* mice, while expression of *Wdfy4* and *Ccr7* did not differ between genotype (fig. S10E). Altogether, these results suggest that loss of CD74 may be associated with transcriptional rewiring of DCs towards a cross-presentation molecular phenotype, prominently involving AP-1 complex members.

### CD74 regulates DC migration and affects CCR7 expression in DCs

The scRNA-seq analysis, using spleen samples of naïve and B16-OVA-bearing mice, revealed differential expression of genes involved in NF-κB and AP-1 signaling pathways. Both transcription factors have previously been implicated in regulating the transcription of *Ccr7* (36). Moreover, given the early accumulation of migratory DCs in the tdLN of *Cd74-/-* mice, we next asked whether CD74 directly regulates DC migration and trafficking by modulating expression of key homing receptors, like CCR7, in steady-state and melanoma (37). At steady state, a significantly higher percentage of CCR7+ MHC-II^high^CD11c^int^ migratory DCs was observed in LNs of *Cd74-/-* mice compared to wild-type controls (Fig. 7A, fig. S11A). Even though there was no difference in *Ccr7* mRNA levels in DCs in the spleen and tdLNs of the B16-OVA-bearing *Cd74-/-* and *Cd74+/+* mice, we assessed CCR7 surface expression in MHC-II^high^CD11c^int^ migratory DCs recruited to the tdLN at different stages of tumor growth, since mRNA and protein levels can significantly differ. Both at the early timepoint and endpoint, *Cd74-/-* MHC-II^high^CD11c^int^ migratory DCs exhibited increased frequency of CCR7+ cells compared to *Cd74+/+* mice (Fig. 7B). These data suggest that CD74 may regulate CCR7 expression during the course of DC migration; however, this regulation likely occurs at a post-transcriptional or post-translational level.

**Figure 7.**
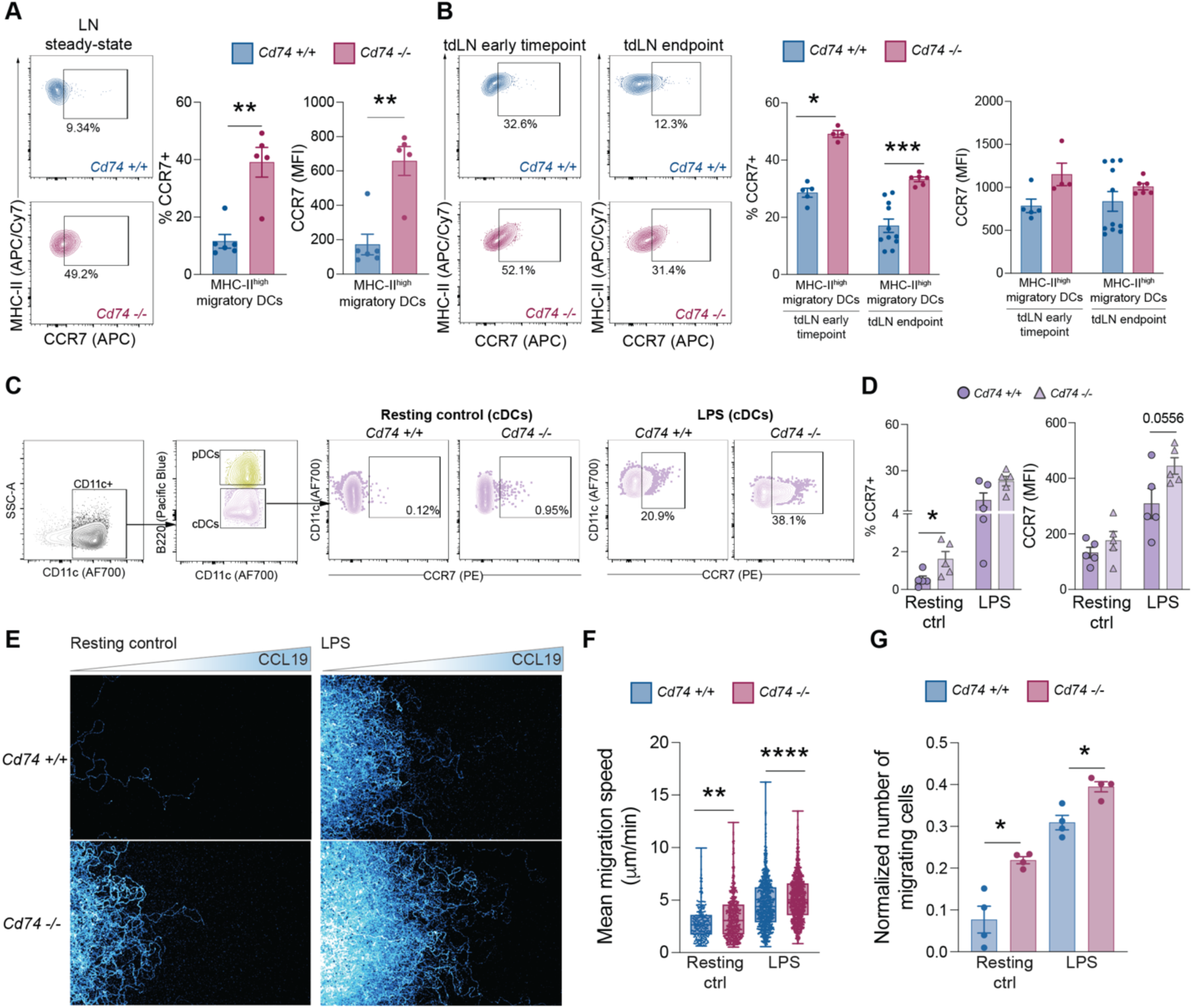
CD74-deficiency results in increased CCR7 expression and enhanced DC chemotaxis. (**A**) Representative flow cytometry plot (left) and frequency of CCR7+ MHC-II^high^ migratory DCs in the lymph node of naïve mice (n=5-6 mice per group, from three independent experiments, mean ± SEM, Mann-Whitney U test). (B) Representative flow cytometry plots (left) and frequency of CCR7+ MHC-II^high^ migratory DCs in the lymph node of B16-OVA-bearing mice at early timepoint and endpoint (n=4-11 mice per group, from two independent experiments, mean ± SEM, Mann-Whitney U test). (**C**) Gating strategy used for the discrimination of different DC populations derived from mFLT3L generated DCs from the bone marrow. Representative flow cytometry contour plots indicating changes in the frequency of CCR7+ cDCs under resting and LPS stimulation conditions. (**D**) Frequency (left) and MFI (right) of CCR7+ cells within cDCs after stimulation with LPS, and resting control (n=5 per group, data points indicate FL-DC cultures from individual mice, from five independent experiments, mean ± SEM, Mann-Whitney U test). (**E**) Representative images of maximum projection of nuclear stain of resting and LPS-matured FL-DCs migrating toward CCL19 (on the right-hand side of the figure panels). (**F**) Mean migration speed of resting and LPS-matured *Cd74+/+* or *Cd74-/-* FL-DCs (data from four independent experiments, cells analyzed: n=539 resting *Cd74+/+*, n=234 resting *Cd74-/-*, n=1110 LPS-matured *Cd74+/+*, n=864 LPS-matured *Cd74-/-*, Mann-Whitney U test) (**G**) Normalized number of migrating cells (data from four independent experiments, mean ± SEM, Mann-Whitney U test). (*\*p≤0.05, **p≤0.01, ***p≤0.001, ****p≤ 0.0001*)

To further explore DC migration, we generated in vitro bone marrow-derived dendritic cells, which resemble steady-state cDCs and pDCs, using FLT3L from *Cd74+/+* and *Cd74-/-* mice. The FL-DC cultures were either resting or matured using LPS or pI:C. Flow cytometry analysis revealed that resting *Cd74-/-* FL-DCs displayed higher percentage of CCR7+ cells compared to *Cd74+/+* cDCs (Fig. 7C, D, fig. S11B). Upon LPS maturation, the frequency of CCR7+ CD11c+B220- FL-DCs (cDC-like cells) and their expression level of the receptor showed an increasing trend (Fig. 7D). pI:C-matured *Cd74-/-* cDC-like cells also showed a tendency toward higher percentages of CCR7+ cells and higher CCR7 expression levels compared to their *Cd74+/+* counterparts (fig. S11D). Resting or LPS-matured CD11c+B220+ FL-DCs (pDC-like cells), on the other hand, did not demonstrate any difference in their CCR7 expression (fig. S11C, D). Therefore, CD74 deficiency enhances CCR7 expression across selected DC subsets both in resting and maturation conditions. This increase in CCR7 levels in *Cd74-/-* FL-DCs aligns well with our in vivo data, further supporting the possibility that CD74 may modulate DC migration capacity.

To determine the functional consequences of increased CCR7 expression on DC migration, we conducted an under-agarose migration assay towards CCL19, a key CCR7 ligand secreted by cells of the lymph nodes, using resting and LPS-matured *Cd74-/-* and *Cd74+/+* FL-DCs (Fig. 7E).

Quantification of migration parameters demonstrated increased mean migration speed (Fig. 7F) and a higher number of migrating cells (Fig. 7G) in *Cd74-/-* FL-DCs. These findings suggest that CD74 impairs DCs migration potentially through regulation of CCR7 expression.

## DISCUSSION

Despite the success of immunotherapy in managing metastatic cutaneous melanoma, therapeutic resistance remains a prevalent challenge in ICB, highlighting the need to better understand mechanisms underlying tumor immune deficiency (1). Current gaps in our knowledge include determining whether the systemic loss of CD74 would ultimately impair or enhance melanoma immunity, given its broad expression across immune cells and its opposing roles in triggering adaptive responses through MHC-II assembly or inducing immune suppression (12,21,38). The phenotypic changes observed in this study resembled those reported in MHC-II-deficient mouse models (39–41). Nonetheless, our results show that CD74 deficiency reprograms cDC subsets into a cross-presenting and pro-migratory state enhancing melanoma immunosurveillance that ultimately augmented CD8+ T-cell priming, infiltration, and effector functions. Therefore, these findings indicate that CD74 shapes melanoma immunogenicity not only through MHC-II-dependent mechanisms but also by modulating cytotoxic T cell-mediated immunity in an MHC-II-independent manner. The involvement of interleukin-1 and NF-κB signaling, revealed by the scRNA-seq analysis, provides a possible mechanistic explanation, indicating that CD74 exerts dual functions as a chaperone and as an immune signaling mediator that modulates tumor progression.

CD74 regulated cDC1 subsets activation and migration in both homeostasis and cancer leading to strengthened adaptive immunity. cDC1s are the initiators of CD8+ T cell-mediated tumor immunity due to their capacity to migrate from the tumor site into tdLNs and cross-present intact tumor antigens via MHC-I to naïve CD8+ T cells (42). In response to IFN-γ and type I interferons, cDC1 in the TME also represent major producers of CXCL10, a chemokine required for effector CD8+ T cell recruitment (43). Here, we observed an increase of CCL5 and CXCL10 in the TME of *Cd74-/-* mice, which could contribute to the increased infiltration of effector CD8+ T cells. Previous studies have demonstrated that CXCL10 is strongly associated with presence of tumor-infiltrating lymphocytes, Th1 immunity, and favorable response to immunotherapy in melanoma (44). Similarly, constitutive CCL5 expression by tumor cells has been implicated in the recruitment of tumor-infiltrating lymphocytes in ovarian cancer (45). Importantly, CCL5-driven recruitment of T cells to the TME can initiate a cascade in which T cells become activated by tumor antigen recognition and promote IFN-γ release, leading to DC activation and CXCL10 secretion, thereby amplifying local T cell infiltration (45). These findings suggest that the chemokine alterations observed in the TME of *Cd74-/-* mice may cooperate with DC mechanisms to further enhance or sustain T cell recruitment.

In addition, we observed that CD74 loss amplifies the DC-driven adaptive effector phase of tumor immunity at several sequential levels. We found an accumulation of activated PD-1+ CD8+ T cells that progress toward exhaustion in the TME, which is a known compensatory feature of inflamed tumors (46), and critical for ICB outcomes (47). Migratory PD-L1+ cDC1s, associated with antigen uptake (48), were also increased in the TME of *Cd74-/-* mice. Although PD-L1 may dampen PD-1+ T cell function, it also protects DCs from CD8+ cytotoxicity, favoring tumor immunity and ICB responses (48). Moreover, the central memory phenotype of CD8+ T cells we observed in *Cd74*-/- mice may also support T cell longevity and rapid activation upon antigen re-exposure, which is critical for durable immunity and ICB responses (49). The enhanced number of IFN-γ+ CD8 T cells in the TME of *Cd74-/-* mice could boost the cytotoxicity of CD8 T cells via autocrine control of granzyme B expression and paracrine induction of CD4 Th1 differentiation (50–53). Ultimately, the increased infiltration of PD-1+ CD8+ T cells resulting from CD74 deficiency translated into significantly enhanced therapeutic responses to PD-1 blockade across two melanoma models. This underscores a critical role for CD74 in restraining effector T cell responses, the loss of which sensitizes tumors to ICB-mediated control.

We also observed an accumulation of pDCs in the TME of *Cd74-/-* mice at the endpoint. Beyond IFN-α secretion after TLR activation, pDCs also exhibit antigen-presentation capacities that can induce immunogenic T-cell responses (54). In mice, pDCs that are activated in vivo by mouse cytomegalovirus induce Th1 polarization, and CpG-activated mouse pDCs expand naïve CD8+ T cells in vitro in response to endogenous and exogenous antigens (55–58). In cancer, pDCs can promote immunogenic anti-tumor responses when appropriately stimulated. In metastatic melanoma patients, vaccination with activated pDCs loaded with tumor-associated peptides elicits favorable CD4 and CD8 T cell responses, while in the B16 melanoma mouse model, TLR-stimulated pDCs mediate tumor killing via TRAIL and GrB as well as through NK-cell activation (59–61). However, tumor pDCs are often in a nonactivated state expressing low CD80/CD86 levels (62). Here, the pDCs in the TME of *Cd74-/-* mice exhibited increased CD80 and MHC-I levels, suggesting that they may participate in local activation of intratumoral T cells. Our data also revealed a shift from lower to higher pDC frequency in the tdLNs of *Cd74-/-* from early timepoint to endpoint. CXCR3-dependent accumulation of circulating pDCs in lymph nodes occurs after exposure to inflammatory stimuli (63). Whether a similar accumulation of pDCs occurs in the tdLNs of *Cd74-/-* mice remains unclear.

This study employed a constitutive *Cd74-/-* mouse model as a proof-of-concept to assess how systemic CD74 depletion reshapes DC-mediated anti-tumor immunity and responsiveness to PD-1 blockade. Use of constitutive *Cd74-/-* permits evaluation of integrated immune effects that might more closely reflect therapeutic targeting of CD74. Moreover, complementary in vitro validation using FL-DCs supports attribution of the observed effects to dendritic cells and their specific subsets. A limitation of this study is that DEA could not be performed on the lymph node samples of the naïve and B16-OVA-bearing *Cd74+/+* and *Cd74-/-* mice due to insufficient DC numbers. Hence, we could not assess whether CD74 deficiency alters DC-T cell interactions in the tdLNs at transcriptional level. Moreover, the discrepancy between the frequencies and absolute numbers of cells in naïve lymph nodes, tdLNs, and the TME could be due to the fact that certain immune populations, like the CD4+ T cells, are dramatically reduced in absolute numbers in the *Cd74-/-*mice, as reported by us and others (22,23). For instance, the dramatic reduction of these immune cells in the naïve and tumor-draining lymph nodes may lead to an increased proportion of other cell populations, such as migratory cDC1s, revealing changes in lymph node composition that may result from altered cellular trafficking.

CD74 has been implicated in the non-canonical endolysosomal MHC-I cross-presentation pathway (64). Using a viral infection model, Basha et al. concluded that CD74 is essential for antigen cross-presentation by directing MHC-I trafficking from the endoplasmic reticulum (ER) to endolysosomes. In contrast, we found that CD74 deficiency enhanced tumor-cell associated antigen cross-presentation and CD8 T cell priming. This discrepancy likely reflects model-specific differences (viral as opposed to tumor antigens). MHC-I typically assembles in the ER with cytosolic peptides; however, it can also traffic through the Golgi apparatus, plasma membrane, and endolysosomes (65). For endosomal viruses that require vacuolar processing and non-cytosolic cross-presentation, CD74 deficiency may restrict MHC-I delivery to endolysosomes, reducing viral antigen-specific CD8+ T cell priming. Nonetheless, in tumors cross-priming mainly follows the cytosolic pathway, with peptides loaded onto MHC-I in the ER (66). In our study, loss of CD74 may favor MHC-I retention in the ER, enhancing peptide loading and surface expression, consistent with the elevated MHC-I levels observed in cDC1s in the TME and tdLNs of *Cd74-/-*mice. Moreover, the increased levels of IFN-α in the TME of *Cd74*-/- mice, likely derived from increased numbers of pDCs, may further support MHC-I upregulation (67). In addition, methodological differences between the two studies, such as DC generation protocols and their maturation states, may also contribute to different results. GM-CSF generated bone marrow-derived DCs (BMDCs) used by Basha et al. resemble inflammatory monocyte-derived DCs, whereas FLT3L promotes cDCs and pDCs development similar to steady-state DCs found in vivo, with the ability to elicit antigen cross-presentation (68,69). Furthermore Basha et al. utilized viral antigen or lethally irradiated ovalbumin-pulsed DCs as source of cell-associated antigens, whereas we employed tumor cell-associated antigens derived by apoptotic B16-OVA melanoma cells, which mimic the antigens found in the tumor microenvironment. Our findings underscore the need to thoroughly understand the role of CD74 in antigen cross-presentation in different biological scenarios before drawing conclusions on its pro- or anti-inflammatory function.

Beyond antigen presentation, our tdLN phenotyping data also revealed a second potential layer of CD74-dependent regulation concerning DC migration. The increased frequency of migratory cDC1s in early timepoint in the tdLNs of *Cd74-/-* mice suggests that CD74 deficiency may promote DC migration in the context of melanoma immunogenicity. Indeed, previous study demonstrated that CD74 is a negative regulator of DC motility using GM-CSF-generated BMDCs stimulated with LPS (17). The study revealed that retention of CD74 caused by knocking out Cathepsin S in DCs disrupts myosin II function by trapping uncleaved CD74 in endolysosomal compartments (17). Therefore, we tested whether CD74 deficiency could improve DC migration using FL-DCs, instead of GM-CSF-generated BMDCs, and implemented an under-agarose migration assay toward CCL19 (70) for single cell tracking. We discovered that *Cd74-/-* FL-DCs migrated more efficiently toward CCL19 compared to *Cd74+/+* controls under both resting and LPS-matured conditions. The previous study investigated how CD74 impacts DC migration only after LPS-induced maturation and attributed their findings to effects on the cytoskeleton. Here, we identified that increase in CCR7 expression and proportion of CCR7+ cells could be a potential driver of the enhanced migratory capacity of *Cd74-/-* FL-DCs, potentially facilitating more efficient antigen delivery to lymph nodes and leading to improved T cell priming. Moreover, the proportion of CCR7+ migratory DCs was consistently increased on in naïve lymph nodes and tdLNs of *Cd74-/-* mice, making the association between CD74 and CCR7 even stronger.

In conclusion, this study identifies CD74 as an emerging checkpoint in DC biology, influencing both antigen cross-presentation and migration to sustain immune responses against melanoma. Future studies phenotyping the TME and secondary lymphoid organs of *Cd74-/-* mice treated with ICB will be necessary to elucidate the mechanisms driving tumor control in a therapeutic context. Moreover, preclinical therapeutic approaches targeting CD74 alone and/or in combinations with ICB will be essential to further establish the impact of CD74 on melanoma immunogenicity and therapeutic potential. Similar efforts are underway for other innate immune regulators to improve tumor immunogenicity (1). Yet, CD74 emerges as a particularly attractive target given its predominant expression in myeloid and B cells and its defined role in DC immunogenicity, unlike broadly expressed enzymes or nuclear regulators. Altogether, our findings position CD74 as a regulator of immunosurveillance and promising candidate for future therapeutic studies aiming to enhance DC-driven immunogenicity in “cold” tumors and improve ICB responses.

## MATERIALS AND METHODS

### Study design

The objective of this study was to investigate the role of CD74 in melanoma progression and its ability in shaping the immune microenvironment of melanoma. For this purpose, we used a constitutive knockout mouse model for *Cd74* and subcutaneously inoculated syngeneic melanoma cell line B16-OVA, which express *Cd74*. To dissect the anti-tumor immune response in the absence of CD74 we conducted spectral flow cytometry analyses with T and myeloid cells in TME and tdLNs. To elucidate the specific role of CD74 in dendritic cells, in vivo cancer cell-associated antigen cross-presentation assay and under-agarose migration studies were performed using unstimulated and LPS-stimulated mFLT3L-generated bone marrow-derived dendritic cells (FL-DCs). To understand how the immune response is influenced by the absence of CD74 in steady-state versus inflammation, we implemented scRNA-sequencing and analyses of immune-related molecular pathways.

### Mice

All mice were maintained and bred in pathogen-free facility according to the guidelines of Central Animal Laboratory (CAL), University of Turku. All animal experiments were conducted at CAL, adhering to the 3Rs and the Finnish Act (62/2006) and approved by the ethical committee (License: ESAVI/39072/2021). Wild-type C57BL/6J and CD74-deficient B6.129S-*Cd74tm1^Liz/J^*mice were purchased from the Jackson Laboratory. C57BL/6-Tg(TcraTcrb)1100Mb/J (OT-I) mice having transgenic OVA-specific CD8+ T-cells were purchased from Charles River Laboratories. All experiments were conducted with 6–10-week-old mice that were age and sex matched.

### Genotyping

Genotyping primers for B6.129S-*Cd74tm1^Liz/J^* were as follows: a) CAC TCA AGG CAA CCT TCC TG (wild-type reverse), b) ATT CGC CAA TGA CAA GAC G (mutant reverse), c) CGC CTG TGG GAA AAA CTA GA (common). The primers were ordered from Integrated DNA Technologies (Integrated DNA Technologies, Inc. (IDT).

### Cell culture

The B16-OVA melanoma cell line (B16F10 melanoma cells stably expressing chicken OVA) was a gift from professor Sirpa Jalkanen. B16-OVA and B16F10 (ATCC, CRL-6475 ™) cells were cultured in complete DMEM (Gibco, 21969-035) supplemented with 10% heat-inactivated Newborn Calf Serum (NBCS) (Gibco, 16010-159), 1% Penicillin/Streptomycin (pen/strep) (Gibco, 15140-122), and 4 mM of ultraglutamine (Lonza, LZBE17-605EU1) at 37°C and 5% CO2. YUMM1.G1 (ATCC, CRL-3363 ™) were cultured in DMEM/F12 (30-2006, ATCC) supplemented with 10% NBCS, 1% NEAA (11440-076, Gibco) and 1% pen/strep at 37°C in 5% CO2.

### Subcutaneous tumor inoculation and treatments with anti-PD1

10^5^ cells were resuspended in 100 μL of sterile phosphate-buffered saline (PBS) (Gibco, 10010-015) and were subcutaneously injected in the right flank of 6–10-week-old male mice. The tumor-bearing *Cd74-/-* and *Cd74+/+* were sacrificed either on day 14 (early time point) or day 20 (endpoint). For selected experiments (Ki-67 expression and CCR7 expression in vivo) female mice were inoculated with 2 x10^5^ B16-OVA for 7 (early time point) or 14 days (endpoint).

For anti-PD1 experiments, mice were subcutaneously injected in the right flank with 1×10⁶ YUMM1.G1 cells or 5×10⁵ B16F10 cells. Once tumor reached a palpable size (50–100 mm³), mice were randomized and treated as follows: mice received intraperitoneal injections of 200μg monoclonal anti–PD-1 antibody (clone J43, BP0033, Bio X Cell) or isotype control (polyclonal Armenian hamster IgG, BE0091, Bio X Cell) for three or four doses at the indicated time points. Mice were sacrificed when tumor volumes exceeded 1,500 mm³.

### Bone marrow-derived dendritic cells preparation with FLT3L (FL-DCs)

Femora and tibiae from mice were flushed with RPMI (Gibco, 61870-010,). Red blood cells were removed with ACK Lysing Buffer (Lonza, BP10-548E). The bone marrow cells were then cultured at a density of 2x10^6^/mL in complete RPMI supplemented with 10% heat-inactivated NBCS, 1% pen/strep, 1X Non-Essential Amino Acids Solution (Gibco, 11140050), 1mM sodium pyruvate (Gibco, 11360070), 10 mM HEPES (Gibco, 15630080), and 0.06 mM 2-Mercaptoethanol (Gibco, 21985023) in the presence of 150 ng/mL recombinant mFLT3L (SinoBiologicals, 51113-M02H) at 37°C and 5% CO2 for 8 days. For the maturation of FL-DCs, loosely adherent cells from the cultured were harvested and incubated with either 100 ng/mL LPS (Sigma, L4516) for 24 hours or 50 μg/mL polyinosinic:polycytidylic acid (pI:C) (InVivoGen, vac-pic) for 24 hours, both followed by a 1-hour of resting in 10% RPMI. The matured FL-DCs and resting controls (no LPS or pI:C) were then used for the agarose migration assay or the analyses of CCR7 cell surface levels.

### Single-cell suspension preparation

Mice were euthanized and organs were immediately harvested. Dissected tumors were cut into small pieces and digested in RPMI supplemented with Liberase TL (Roche, 5401020001) and DNase I (Sigma Aldrich, D5025-15KU) at 37°C for 30 min with shaking. RPMI supplemented with 2% NBCS was added in the digested tissues. These were then passed through a sterile cell strainer with 70 µm nylon mesh (FisherBrand, 22363548). The samples were washed and the pellets were resuspended with PBS and were purified on a Histopaque-1077 (Sigma Aldrich, 1077-500 mL) discontinuous gradient centrifugation at 2000 rpm for 20 min at room temperature without brake. The collected buffy coats were then washed and the cell pellets were quantified using dual-chamber cell counting slides (Bio-Rad, 1450011) and a TC20 automated cell counter (Bio-Rad, 1450102).

Lymph nodes were cleaned from the fat tissue. After mechanical dissociation, they were enzymatically digested using 1 mL RPMI supplemented with Liberase TL (Roche, 5401020001) and DNase I (Sigma Aldrich, D5025-15KU) at 37°C for 20 min. RPMI supplemented with 2% NBCS was used to stop the digestion and the tissue was passed through a sterile cell strainer with 70 µm nylon mesh. The cell numbers were then determined using the same method as described for the tumor tissue.

### Flow cytometry

Cells were washed with 1X PBS and centrifuged at 300 x g for 5 min. Cell pellets were first incubated with either yellow or red fixable viability dye depending on the experiment. After a washing step, cells were incubated with anti-mouse CD16/32 for 10 min in 4°C and then incubated with an antibody cocktail. The antibodies and reagents used for flow cytometry are listed in Table S1. The antibodies were diluted in flow cytometry staining buffer (2% BSA, 0.5 mM EDTA, in PBS). After the antibody incubation, cells were washed with flow cytometry staining buffer and the data were acquired on a 3-laser Northern Lights spectral flow cytometer (CyTEK Biosciences). The spectral unmixing was performed using SpectroFlo (CyTEK Biosciences), and the data were further analyzed with FlowJo software (version 10.10.0).

For staining with the OVA peptides, blood samples were collected in EDTA tubes (BD, 365975), and incubated with SIINFEKL MHC Dextramer (Immudex, JD02163) or negative control peptide (SSYSYSSL) MHC Dextramer (Immudex, JD03553) in staining buffer (PBS, 1 mM EDTA, 2% NBCS, pH 7.4) at RT in the dark, for 10 min. Without washing the samples were blocked with anti-mouse CD16/32 at RT for 10 min followed by incubation with relevant antibodies in 4°C for 20 min (Table S1). Red blood cells were lysed with 1X BD FACS™ Lysing Solution (BD, 349202) for 10 min at room temperature. Samples were washed twice with staining buffer and resuspended in 150 μL for acquisition on a 3-laser Northern Lights spectral flow cytometer.

For cytokine assessment, 10⁶ cells were stimulated with 25 ng/mL phorbol 12-myristate 13-acetate (Sigma-Aldrich, P8139) and 1 μg/mL ionomycin (Sigma-Aldrich, I9657) in the presence of Brefeldin A (BioLegend, 420601) for 4 h. After surface staining, cells were permeabilized with the BD Cytofix/Cytoperm™ Kit (BD, 554714) following the manufacturer’s instructions. Cells were incubated with Fixation/Permeabilization solution for 30 min at RT in the dark, washed with 1X BD Perm/Wash Buffer, and centrifuged. Pellets were then incubated with antibodies in 1X Perm/Wash Buffer for 30 min at RT in the dark, washed twice, and resuspended in flow cytometry staining buffer. For intracellular staining of Foxp3 and Ki-67, cells were permeabilized after surface marker staining using the Foxp3/Transcription Factor Staining Buffer Set (Invitrogen, 00-5523). Cells were resuspended in the residual volume, incubated with antibodies for 30 min at RT in the dark, washed twice with Permeabilization Buffer, and finally resuspended in ion-free PBS containing 1 mM EDTA.

### Intratumoral cytokine quantification

Tumor tissue was homogenized on ice in Tissue Protein Extraction Reagent (Thermo Scientific, #78510) supplemented with Protease/Phosphatase Inhibitor Cocktail (Cell Signaling, 5872S). Soluble lysates were centrifuged at 5,000 × g for 5 min at 4 °C, and cytokine concentrations were normalized to tumor weight. Intratumoral cytokine quantification was performed via flow cytometry and the LEGENDplex™ Mouse Anti-Virus Response Panel (Table S1) according to manufacturer’s instructions. Data acquisition was done using a 3-laser Northern Lights spectral flow cytometer and data analysis was performed with BioLegend’s LEGENDplex™ data analysis software.

### Under-agarose migration assay

The under-agarose migration assay was performed as previously described (70). Briefly, 1% UltraPure Agarose (Invitrogen, 16500100) was mixed in phenol red-free RPMI medium with 1x Hanks’ balanced salt solution (pH 7.3; Sigma Aldrich, H1387) and 10 % FBS. The agarose-containing medium was poured onto custom-made chambers with a diameter of 12 mm and allowed to polymerize for 35 min. Two holes of 2 mm diameter were then punched into the agarose 3 mm apart, and either 5x10^5^ FL-DCs or 17.5 ng of CCL19 in phenol red-free RPMI were added into the holes. FL-DCs were stained with NucBlue (Invitrogen, R37605) for 15 min at 37°C and washed once with RPMI before being injected into the agarose hole. Cell migration was recorded with an inverted Nikon Eclipse Ti2-E widefield fluorescent microscope at 37°C with 5 % CO2 for 8 hours with 1-minute intervals and 4x objective.

### Image analysis and statistics

Fiji was used for image processing and analysis, and all graphs and statistical analyses were performed with GraphPad Prism.

### Quantification of cell migration

Cell migration was quantified from 2200x1500 μm size region-of-interests positioned next to the cell source. Migration of Nucblue (Invitrogen, R37605) labeled cells was analyzed using TrackMate plugin for automated cell tracking with carefully adjusted parameters and filtering to exclude unmoving dead cells and fragmented tracks (track displacement > 60 μm, track duration > 120 min). The same parameters were applied consistently in different stimulation conditions among the samples to obtain comparable results.

Maximum projections of the Nucblue channels were created in Fiji from the same ROIs, and Cyan Hot lookup table was added to the images after smoothing and background subtraction.

### In vivo cell-associated antigen cross-presentation assay

On day 0, 70-80% confluent B16-OVA melanoma cells were incubated with 1 μg/mL (1.8 μM) of doxorubicin hydrochloride (Sigma Aldrich, 44583-1MG) to induce apoptosis. On the same day, spleens and inguinal lymph nodes were harvested from OT-I mice. Following tissue dissociation, the naïve CD8+ T cells were enriched using Magnetic Cell Separation and the CD8a+ T cells isolation kit for mouse (130-104-075, Miltenyi Biotec). The enriched naïve OT-I CD8+ T cells were then labeled with 1 μM of CellTrace™ Violet Cell (CTV) (ThermoFisher, C34571) according to manufacturer’s protocol. Subsequently, 2.5 x 10^6^ CTV-labeled OT-I CD8+ T cells were intravenously injected into *Cd74-/-* and *Cd74+/+* recipient mice. On day 1, the doxorubicin-treated (apoptotic) B16-OVA melanoma cells were harvested and 1 x 10^6^ cells were subcutaneously injected on the right flank of each recipient mouse. After 5 days, blood, tumor-draining lymph nodes (right inguinal LN), and spleens were collected. Single-cell suspensions were prepared from secondary lymphoid organs. Blood samples were stained using the protocol described above, and OT-I CD8+ T cell proliferation was assessed by CTV dilution within the CD8+ SIINFEKL Dextramer+ population. Single-cell suspensions of the tdLNs and spleens were stained using LIVE/DEAD Yellow viability dye, anti-CD45, anti-CD8, anti-TCRb5.1 5.2, and anti-CD44, as detailed in Table S1. Proliferation was quantified using flow cytometry based on CTV dilution within the CD44^high^ CTV+ population of OT-I CD8+ T cells.

### In vitro antigen presentation assay

FL-DCs were generated as described above. On day 8-9 of differentiation, loosely adherent FL-DCs were harvested and pulsed for 24h with 1 or 2 mg/mL chicken ovalbumin (OVA) (MP biomedicals, 191224), or for 1h at 37°C with varying concentrations of SIINFEKL (OVA 257-264) (Invivogen, vac-sin), the H-2Kb-restricted peptide epitope of ovalbumin, prior to co-culture with OT-I CD8+ T cells. Naïve OT-I CD8⁺ T cells were enriched and labeled with CellTrace™ Violet (CTV) as described above. Following labeling, 2 x 10^5^ OT-I CD8+ T cells were co-cultured with 4 x 10^4^ OVA-pulsed FL-DCs at 1:5 or 1:10 ration (T cells: DCs) in 96-well U bottom plates. After 3 days, co-cultured cells were stained with anti-CD8, Zombie NIR™ Fixable Viability Kit, and anti-CD44 (Table S1) at 4°C for 20 min in the dark. OT-I CD8+ T cell proliferation was evaluated using flow cytometry based on CTV dilution within CD44^high^ population.

### DQ-OVA antigen uptake and proteolysis assays

FL-DCs were generated as described above. Loosely adherent cells from the culture were harvested and incubated with 100 ng/mL LPS (Sigma, L4516) for 24 followed by a 1-hour of resting in 10% RPMI. Then, 2.5 x 10^5^ LPS-matured FL-DCs and resting controls (no LPS) were incubated with 40 µg/mL DQ-OVA (Invitrogen) for 5, 15, 30, and 60 minutes at 37 °C. Moreover, 2.5 x 10^5^ LPS-matured FL-DCs and resting controls were incubated with 40 µg/mL DQ-OVA for 60 minutes at 4°C as controls. The fluorescence produced by the DQ-OVA was measured using flow cytometry and the FITC channel within CD11c+ cells. The frequency of DQ-OVA+ cells and MFI acquired from the samples incubated at 4°C was then subtracted from the rest of the samples.

### Single-cell RNA sequencing (scRNA-seq)

Single-cell suspensions were prepared as described above from *Cd74+/+* and *Cd74-/-* naïve and tumor-draining lymph nodes, as well as from naïve spleens and spleens of tumor-bearing mice. Sample integrity and RNA quality were assessed using the Agilent Bioanalyzer 2100 (Agilent Technologies), and the concentration was measured with Qubit ®/Quant-IT® Fluorometric Quantitation (Life Technologies). All samples passed the quality assessment.

Single-cell RNA-seq libraries were prepared using the Chromium Next GEM Single Cell 3’, v3.1 reagent kit (10XGenomics) according to the manufacturer’s instructions. 10^4^ cells per sample were loaded on the Chromium controller (10XGenomics) to achieve optimal single-cell encapsulation. Dual indexing was performed using the Dual Index Kit TT Set A (10XGenomics).

Library fragment size and concentration were evaluated with Agilent Bioanalyzer 2100 and Qubit®/Quant-IT® Fluorometric Quantitation, respectively. The average library fragment size was 471 bp.

Sequencing was performed at Finnish Functional Genomics Centre (FFGC, Turku Bioscience Centre) on an Illumina NovaSeq 6000 instrument using a S4 v1.5 flow cell. Libraries were sequenced with paired-end sequencing (2x50 bp) and a ≥ 90% Q30 base quality threshold. Base calling and demultiplexing were performed with CellRanger v7.2.0 (10XGenomics) using cellranger mkfastq. Raw data were delivered in FASTQ format.

### Analysis of scRNA-seq data

The detailed methodology used to analyze the scRNA-seq data from spleen and right inguinal lymph node from naïve *Cd74-/-* and *Cd74+/+* mice, as well as from spleen and tumor-draining lymph node from B16-OVA-bearing *Cd74-/-* and *Cd74+/+* mice are described in Supplementary Material and Methods.

### Statistical analyses

GraphPad Prism (GraphPad Software, version 10) was used to perform all statistical analyses. Two-way ANOVA was used to assess if there is significant effect of genotype in tumor growth. The normality of the data was evaluated using the Kolmogorov-Smirnov test. To determine significance, unpaired Student’s t test was performed for normally distributed data and Mann-Whitney test for data that do not follow Gaussian distribution. The number of independent biological replicates and experiments used for the graphs and statistical tests are described in figure legends.

### Supplementary Material Supplementary figures

Figs. S1 to S11

### Supplementary tables

Table S1. Flow cytometry antibodies and related reagents

## Acknowledgments

The scRNA sequencing study was supported by Turku Bioscience Center, Finnish Functional Genomics Centre, University of Turku and Åbo Akademi and Biocenter Finland. We also thank Clara-Theresia Kolehmainen, Guillermo Martinez Nieto, and Petra Sipilä from Turku Center for Disease Modeling for performing the genotyping of the B6.129S-Cd74tm1Liz/J mice.

## Funding

This research was supported by:

Research Council of Finland grants 330876, 336530, 358209 (CRF);

Research Council of Finland InFLAMES Flagship grants 337530, 357910, and 358823;

Research Council of Finland grant 352727 Profiling area "Capitalising Immunity to Combat Disease";

Sigrid Juselius Foundation grants 220022, 230026 (CRF);

Jane and Aatos Erkko Foundation grants 5957-12539 and 7180-2BD08 (CRF); Turun Yliopistosäätiö 081497/320E25 (CRF);

Research Council of Finland 362007 (JA); Instrumentarium Science Foundation (JA);

ImmuDocs National Doctoral Education Pilot Program (PW); Turku Doctoral Programme of Molecular Medicine (EM); InFLAMES Doctoral module (EM);

The Maud Kuistila Memorial Foundation (EM)

## Author contributions

Conceptualization: EM, GP, CRF Methodology: EM, GP, JA, MS, CRF Investigation: EM, GP, PW, JA, OP Visualization: EM, GP, PW, JA, OP

Funding acquisition: CRF Project administration: CRF Supervision: MS, CRF

Writing – original draft: EM, MS, CRF

Writing – review & editing: EM, GP, PW, JA, OP, SEC, MS, CRF

## Data and materials availability

We aim to deposit the scRNA-seq data generated in this study in public repositories (e.g., GEO) upon manuscript acceptance. The codes for reproducing the analysis and figures are available on GitHub (https://github.com/MedicalImmunoOncologyResearchGroup/CD74). Used python and R packages are described in the supplementary materials and methods section.

