## Supplementary Material for "CD74 regulates antitumor immunity in melanoma by reprogramming dendritic cell immunogenicity and migration"

Eleftheria Maranou *et al.*

**The PDF file includes:**

Materials and Methods
Table S1
Fig. S1 to S11

### Materials and Methods

**Table S1.** Antibodies and reagents used for flow cytometry

| Reactivity | Fluorophore | Clone | Cat. No | Company |
|---|---|---|---|---|
| anti-mouse CD8a | FITC | 53-6.7 | 100706 | Biolegend |
| anti-mouse CD62L | PE | MEL-14 | 104408 | Biolegend |
| anti-mouse/human CD44 | PE/ Cyanine 7 | IM7 | 103030 | Biolegend |
| anti-mouse CD25 | APC | PC61 | 102012 | Biolegend |
| anti-mouse CD3 | APC/Cy7 | 17A2 | 100222 | Biolegend |
| anti-mouse CD45 | violetFluor™ 500 | 30-F11 | 85-0451-U025 | CyTEK |
| anti-mouse CD4 | cFluor® V610 | RM4-5 | R7-20270 | CyTEK |
| anti-mouse/human Foxp3 | PerCP/eFluor™ 710 | FJK-16s | 46-5773-82 | eBioscience |
| anti-mouse/human Ki-67 | Brilliant Violet™ 786 | SolA15 | 417-5698-82 | Invitrogen |
| anti-mouse PD-1 | PE/eFluor™ 610 | J43 | 61-9985-82 | eBioscience |
| anti-mouse CD69 | Brilliant Ultra Violet™ 737 | H1.2F3 | 367-0691-82 | |
| anti-mouse/human Granzyme B | APC | QA16A02 | 372204 | Biolegend |
| anti-mouse IFN-γ | PE | XMG1.2 | 505808 | Biolegend |
| anti-mouse Ly6C | PerCP/Cyanine 5.5 | HK1.4 | 128012 | Biolegend |
| anti-mouse CD11c | PE/Cyanine 7 | N418 | 117318 | Biolegend |
| anti-mouse CD3 | PE/Cyanine 5 | 17A2 | 100274 | Biolegend |
| anti-mouse/human CD11b | APC | M1/70 | 101212 | Biolegend |
| anti-mouse/human B220 | redFluor™ 710 | RA3-6B2 | 80-0452-U100 | CyTEK |
| anti-mouse MHC-II | APC/Cyanine 7 | M5/114.15.2 | 107628 | Biolegend |

| | | | | |
|---|---|---|---|---|
| anti-mouse Ly6G | violetFluor™ 450 | 1A8 | 75-1276-U100 | CyTEK |
| anti-mouse CD80 | Brilliant Violet™ 480 | 16-10A1 | 414-0801-82 | eBioscience |
| anti-mouse MHC-I (H-2kb) | Super Bright 702 | AF6-88.5.5.3 | 67-5958-82 | eBioscience |
| anti-mouse CD103 | Brilliant Ultra Violet 737 | 2E7 | 367-1031-82 | eBioscience |
| anti-mouse CD24 | Super Bright 645 | M1/69 | 64-0242-82 | eBioscience |
| anti-mouse/human CD44 | Brilliant Violet 605™ | IM7 | 103047 | Biolegend |
| anti-mouse CD11c | Alexa Fluor® 700 | N418 | 117320 | Biolegend |
| anti-mouse/human B220 | Pacific blue | RA3-6B2 | 103227 | Biolegend |
| anti-mouse CD24 | APC | M1/69 | 101814 | Biolegend |
| anti-mouse CCR7 | APC | 4B12 | 17-1971-82 | eBioscience |
| anti-mouse CCR7 | PE | 4B12 | 120106 | Biolegend |
| anti-mouse TCR Vβ5.1, 5.2 | PE/Cyanine 7 | MR9-4 | 139508 | Biolegend |
| TruStain FcX™ (anti-mouse CD16/32) | n/a | 93 | 101320 | Biolegend |
| MHC-I Dextramer SIINFEKL (H-2Kb) | APC | n/a | JD02163 | Immudex |
| MHC-I Dextramer SSYSYSSL (H-2Kb) Negative control | APC | n/a | JD03553 | Immudex |
| LEGENDplex™ MU Anti-Virus Response Panel | n/a | n/a | 740622 | Biolegend |
| LIVE/DEAD™ Fixable Yellow Dead Cell Stain | 405/570 nm | n/a | L34967 | Invitrogen |
| ViaDye™ Red Fixable Viability Dye | 615/740 nm | n/a | R7-60008 | CyTEK |
| Zombie NIR™ Fixable Viability Kit | 640/743 nm | n/a | 423105 | Biolegend |

**Analysis of scRNA-seq data**

**Data Processing and Quality Control**

Raw sequencing data was processed using the 10x Genomics Cell Ranger pipeline. Initial dataset comprised 8 samples: *Cd74-/-* and *Cd74+/+* tumor-draining lymph node (tdLN), naive lymph node, spleen from tumor-bearing mice, and naive spleen. Quality control metrics included: minimum threshold of 100 genes per cell, mitochondrial gene content < 20%, ribosomal gene content < 50%, hemoglobin gene content < 2%. Cells were filtered based on gene count distribution with lower cutoff at 1st percentile and upper cutoff at 98th percentile. Doublet detection was performed using Scrublet. Expected doublet rates were calculated using a linear model based on cell recovery numbers. Doublet scores were computed for each sample independently, and cells predicted as doublets were removed from downstream analysis.

Detected doublet rates were1.1-3.4% in lymph nodes and 1.5-2.3%. inspleen. Data integration was performed using Scanorama. Highly variable genes (HVGs) were identified using Seurat v3 methodology preserving genes that were highly variable at least in one sample. Integration steps includednormalization to 10,000 counts per cell, log-transformation, PCA computation (30 components), neighbor graph construction using Scanorama-transformed data, UMAP and t-SNE visualization.

**Clustering and Iterative Refinement**

Clustering was performed using the Leiden algorithm (sc.tl.leiden) at multiple resolutions to capture both broad and fine cell populations. Initial clustering was performed at resolution = 1.0 (leiden_initial). Clusters containing DCs, monocytes, and macrophages were further refined by subsetting and re-clustering at lower resolutions (e.g., 0.3 for cDC1 and cDC2 sub-clusters in spleen samples). The final cluster assignment was set to the most refined clustering. Uniform Manifold Approximation and Projection (UMAP) was used to visualize cell populations (sc.pl.umap).

**Cell Type Annotation**

Initial cell type predictions were performed using CellTypist (v1.3.0) with the Immune_All_Low.pkl model. Mouse-to-human gene symbol conversion was conducted using the Jackson Laboratory's HomoloGene mapping file, ensuring compatibility with CellTypist's human reference models. Clusters were annotated based on differential expression analysis (DEA) and canonical marker genes curated from PanglaoDB, Tabula Muris, ACT (Annotation of Cell Types, formerly CellMarker). DEA was performed using the Mann-Whitney U test (i.e. Wilcoxon rank-sum test, sc.tl.rank_genes_groups with method = 'wilcoxon', corr_method='benjamini-hochberg'). Genes with adjusted p-values < 0.03 and log2 fold-change > 1.0 were considered significant. Custom annotation dictionaries were created for both low- and high-resolution cell type assignments, mapping cluster IDs to biological cell types (e.g., "Naive B cells," "Lymphoid-resident cDC1,"). These annotations were stored in the cell_type_low_res and cell_type_high_res columns of the AnnData object.Gene module scores for key immune populations (B cells, T cells, DC subsets, monocytes, macrophages, plasma cells, etc.) were calculated using sc.tl.score_genes. Marker sets were derived from published literature and public databases (29-32). UMAPs displaying module scores were generated to validate cell type assignments and subpopulation identities.

**Differential Gene Expression of *Cd74-/-* and *Cd74+/+* in Various Cell Types**

Differential gene expression between conditions was assessed using the Mann–Whitney U test (two-sided) for each gene within each cell type or aggregated group, requiring at least two cells per group for inclusion. p-values were corrected for multiple testing using the Bonferroni method. For combined or subsetted cell populations (e.g., "Combined DCs"), data from all relevant subtypes were aggregated prior to statistical testing.

**Pseudobulk Differential Gene Expression Analysis with Interactions**

For each spleen sample and cell type group from DC subtypes, raw count matrices were aggregated by summing counts across all cells belonging to each sample-cell type combination. To increase robustness and account for biological variability, each sample was randomly subdivided into four pseudobulk replicates. For each replicate, cells were randomly assigned and counts summed, yielding four pseudobulk samples per condition (naïve/pathogenic, referring to tumor-bearing) per group (*Cd74-/-* and *Cd74+/+*). The resulting pseudobulk count matrices were used as input for downstream DEA. DEA was performed using the PyDESeq2 package, which implements the DESeq2 methodology for bulk RNA-seq adapted to pseudobulk single-cell data. The design formula included genotype, pathogenicity, and their interaction: ~genotype + pathogenicity + genotype:pathogenicity. The interaction analysis was carried out using R version of DESeq2. For each comparison, Wald tests were performed to estimate log2 fold changes and significance, Multiple testing correction was applied using the Benjamini-Hochberg false discovery rate (FDR) method, and a pseudocount (eps = 1e-300) was added for numerical stability in log-transformation of p-values. Genes were considered significantly differentially expressed if they met both log2 fold change and adjusted p-value thresholds $|\log2FC| > 0.5$ and $padj < 0.05$. Volcano plots were constructed to visualize the relationship between log2 fold change and significance (log-transformed adjusted p-values), with top genes labeled directly on the plots. Clustermaps (heatmaps) were generated using log1p-normalized pseudobulk expression and genewise Z-score normalization for the top differentially expressed genes, comparing *Cd74+/+* and *Cd74-/-* across conditions.

**Gene Ontology**

Differentially expressed genes from DC pseudobulk interaction DEA were used for gene ontology analysis (GOA) with python package GOATOOLS. GO terms were filtered for immune relevance by including all GO terms higher in the tree for lists of GO terms determined as core immune processes, DC-specific processes, tumor microenvironment relevant, cellular processes relevant to DCs, and additional immune processes (Data file S3, for a full list of terms), and further by a long list of immune-related keywords. Finally, only GO terms with at least 5% of the pathway genes represented by our list of DEGs, and with p_corrected<0.01 as well as p_corrected<0.03 (FDR: Benjamini-Hochberg), were included. The GO terms were then clustered into clusters of Jaccard similarity greater or equal to 0.4, and the clusters were given descriptive names.

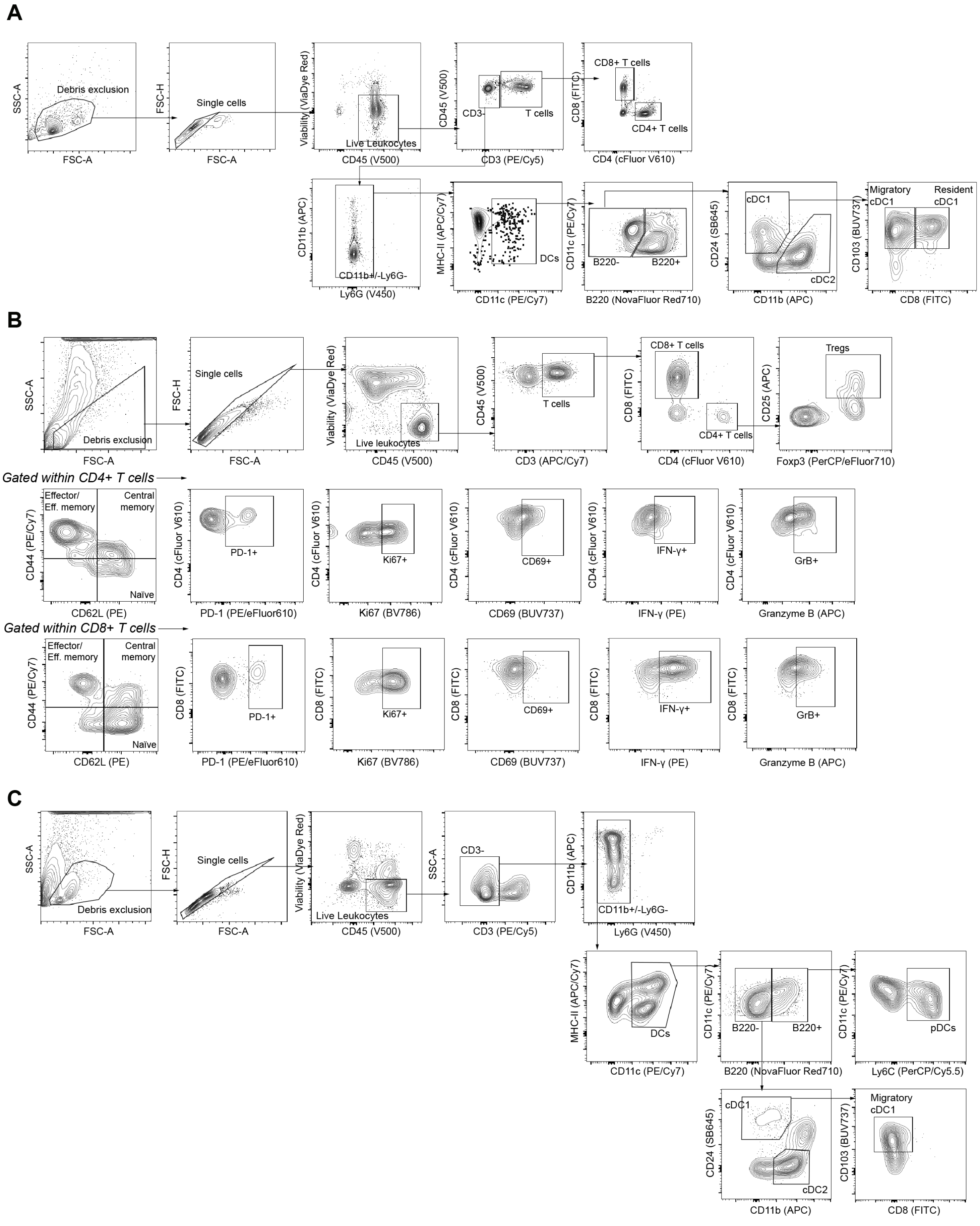


**Fig. S1. Gating strategies for the phenotyping of dendritic cells and T cells in naïve inguinal lymph node, tumor-draining lymph node (tdLN), and tumor microenvironment (TME). A,** Gating strategy followed for phenotyping of dendritic cells and T cells in naïve lymph nodes. **B,** Gating strategy for phenotyping of T cells and their subsets in TME of B16-OVA-bearing mice. **C,** Gating strategy to discriminate dendritic cells and their subsets in tdLN in early timepoint and endpoint of the B16-OVA inoculation experiments.

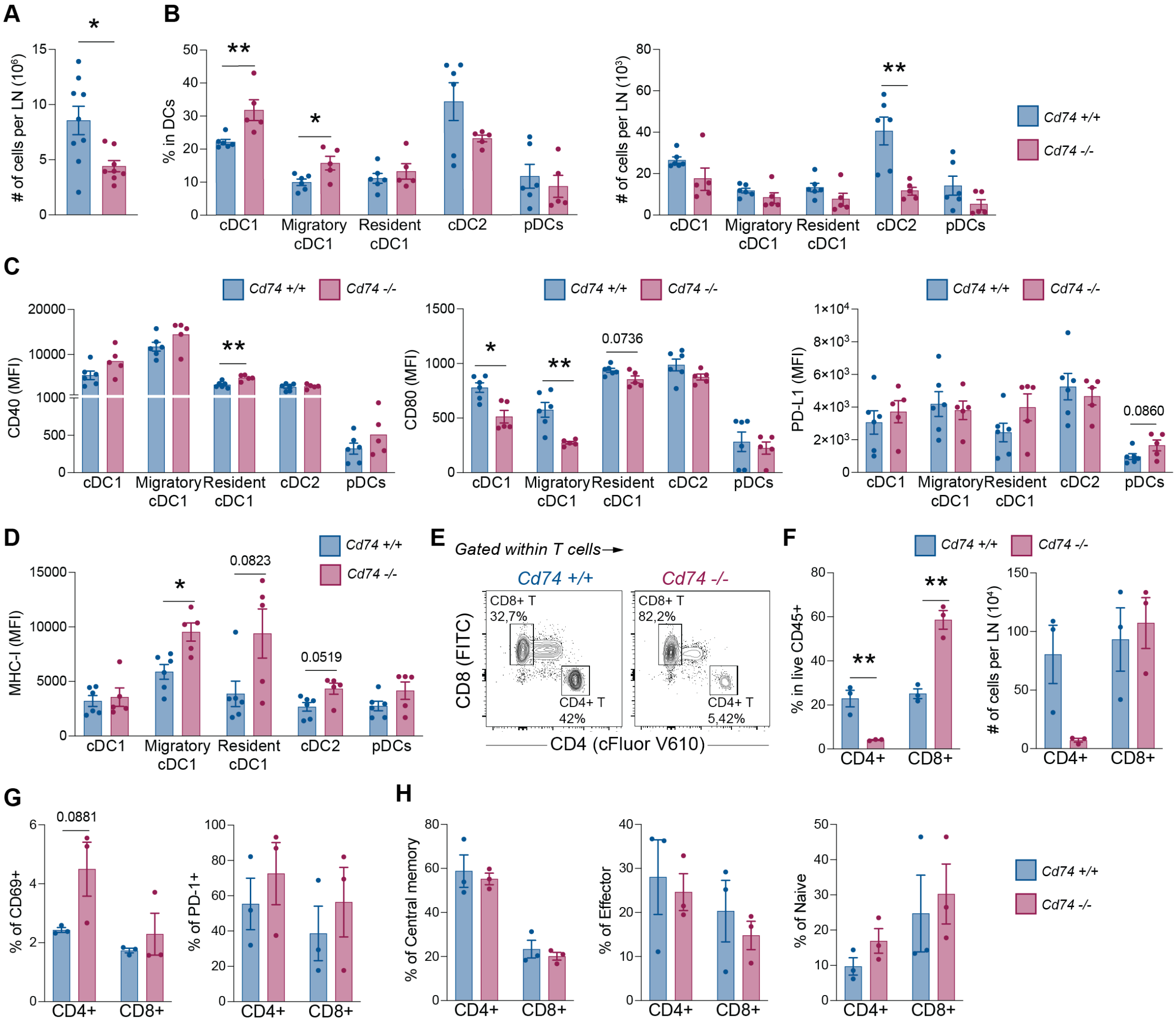


**Fig. S2. Analyses of dendritic cell and T-lymphocyte phenotypes in lymph nodes of *Cd74-/-* mice under homeostasis. A,** Absolute number of cells per naïve inguinal lymph node (n=8-9 mice per group, from four independent experiments, means ± SEM, Mann-Whitney U test). **B,** Frequency (left) and absolute number (right) of DC subsets within DCs in naïve inguinal lymph node (n=5-6 mice per group, from three independent experiments, means ± SEM, unpaired t-test). **C,** Median fluorescence intensity (MFI) of DC-activation molecules CD40, CD80, and PD-L1 in different DC subsets of the naïve inguinal lymph node (n=5-6 mice per group, from three independent experiments, means ± SEM, Mann-Whitney U test). **D,** MFI of MHC-I in different DC subsets of the naïve inguinal lymph node (n=5-6 mice per group, from three independent experiments, means ± SEM, Mann-Whitney U test). **E,** Representative flow cytometry contour plot for the frequency of CD4+ and CD8+ T cells in the naïve inguinal lymph node. **F,** Frequency (left) and absolute number (right) of CD4+ and CD8+ T cells within leukocytes (CD45+) in the naïve inguinal lymph node (n=3 mice per group, from one experiment, means ± SEM, unpaired t-test). **G,** Frequency of early activation marker CD69 (left) and T cell-exhaustion marker PD-1 (right) within CD4+ and CD8+ T cells in the naïve inguinal lymph node (n=3, from one experiment, means ± SEM unpaired t-test). **H,** Frequency of central memory, effector, and naive CD4+ as well as CD8+ T cells in the naïve inguinal lymph node (n=3, from one experiment, means ± SEM, unpaired t-test). (*$p \leq 0.05$, **$p \leq 0.01$, ***$p \leq 0.001$, ****$p \leq 0.0001$)

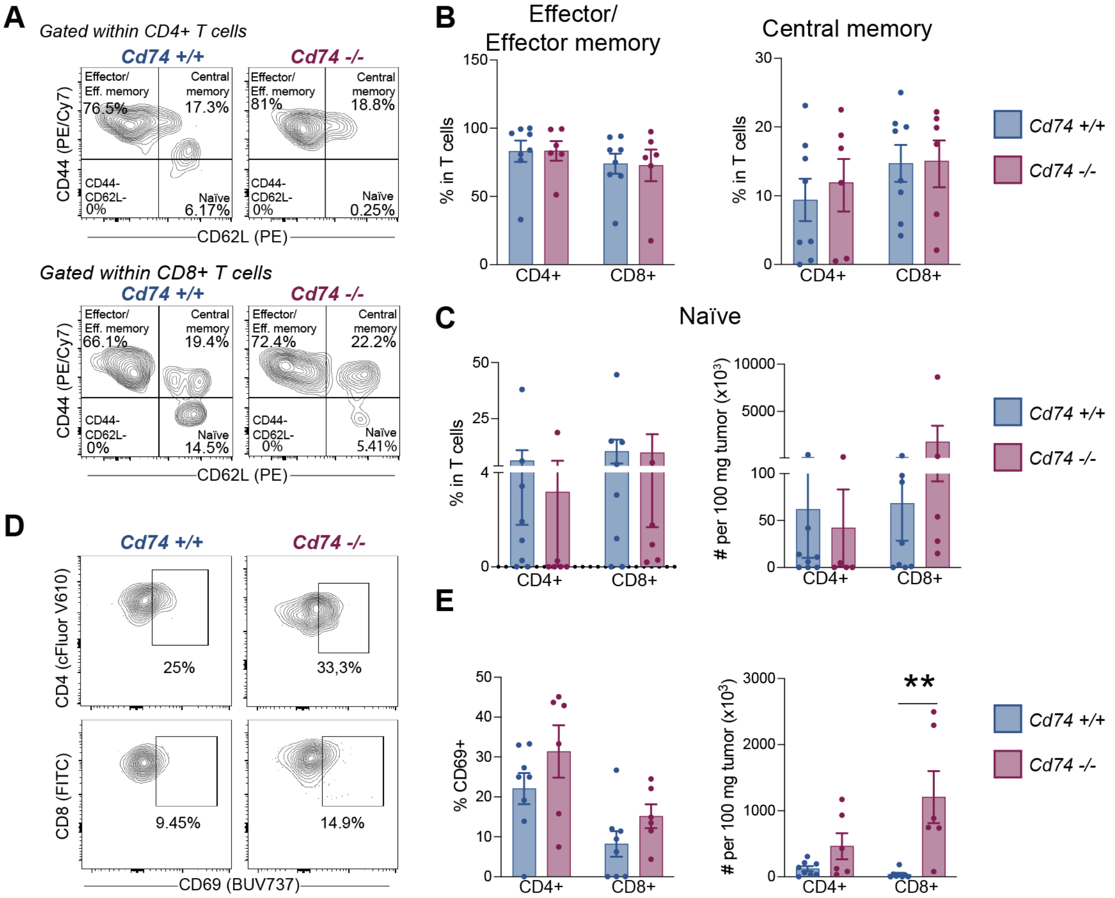


**Fig. S3. Analyses of T-cell phenotypes in in the TME. A,** Representative flow cytometry contour plots for the frequency of CD4+ and CD8+ effector/effector memory, central memory, and naïve T cells in the TME. **B,** frequency of CD4+ and CD8+ effector/effector memory, central memory, and naïve T cells in the TME. (n=6-8 mice per group, from two independent experiments, mean ± SEM, Mann-Whitney U test). **C,** Frequency and total number of naïve CD4+ and CD8+ T cells per 100 mg of tumor tissue (n=6-8 mice per group, from two independent experiments, mean ± SEM, Mann-Whitney U test). **D,** Representative flow cytometry contour plots for the frequency of CD69+ cells within the CD4+ and CD8+ T-cell populations in the TME. **E,** left: Frequency of CD69+ CD4+ and CD69+ CD8+ T cells calculated within the parent populations, CD4+ and CD8+, respectively (n=6-8 mice per group, from two independent experiments, mean ± SEM, Mann-Whitney U test); right: Total number of CD69+ cells within the CD4+ and CD8+ T-cell populations per 100 mg of tumor tissue (n=6-8, from two independent experiments, means ± SEM, Mann-Whitney U test). (**p≤0.05*, ***p≤0.01*, ****p≤0.001*, *****p≤0.0001*)

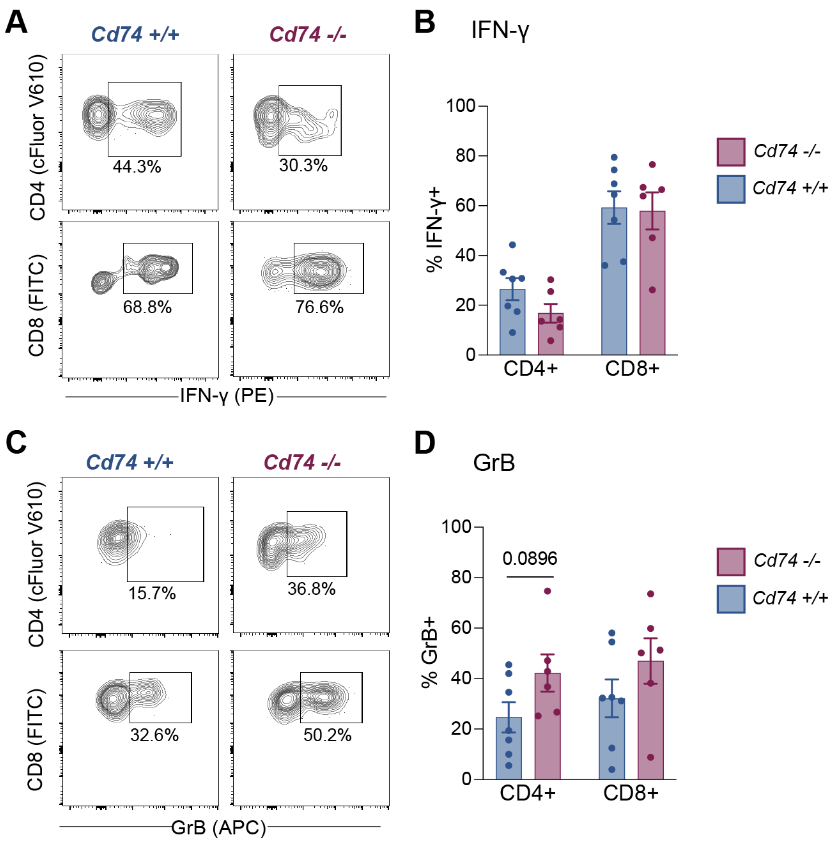


**Fig. S4. Frequencies of IFN-γ and granzyme B in in the TME. (A to D) Representative flow cytometry** contour plots (A and C) for analyzing the frequency of (B) IFN-γ+ and (D) GrB+ cells within the CD4+ and CD8+ T-cell populations in the TME (n=6-7 mice per group, from two independent experiments, means ± SEM, unpaired t-test for frequency, Mann-Whitney U test for total cell amounts). (**p≤0.05*, ***p≤0.01*, ****p≤0.001*, *****p≤0.0001*)

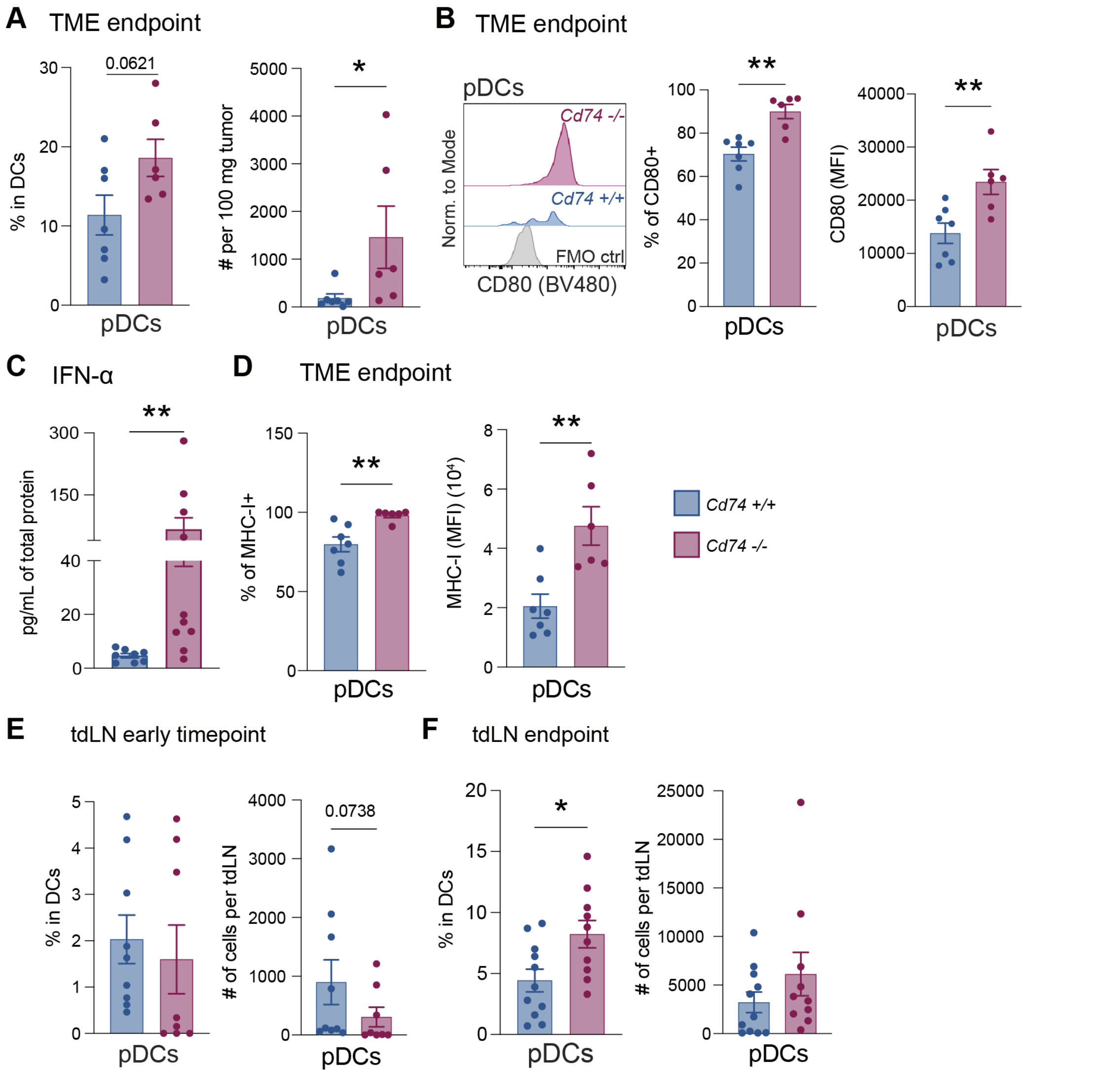


**Fig. S5. Analyses of pDCs in the TME of *Cd74-/-* and *Cd74+/+* mice. A,** Frequency and total number of pDCs per 100 mg of tumor tissue. **B,** Representative histograms for the expression of CD80 on the surface of pDCs, frequency of CD80+ pDCs, and MFI of CD80 calculated within pDCs in the TME (n=6-7 mice per group, from two independent experiments, means ± SEM, unpaired t-test for frequencies and Mann-Whitney U test for cell number and MFI). **C,** Intratumoral IFN-α quantification via LEGENDplex™ in B16-OVA tumors in the endpoint of the tumor challenge experiments (n=8-10 mice per group, from two independent experiments, means ± SEM, Mann-Whitney U test). **D,** Frequency of MHC-I+ pDCs and MFI of MHC-I calculated within pDCs in the TME (n=6-7 mice per group, from two independent experiments, means ± SEM, Mann-Whitney U test). **E,** Frequency (left) and absolute number (right) of pDCs within DCs in the early timepoint tdLNs post B16OVA sc challenge (n=8-9 mice per group, from two independent experiments, mean ± SEM, Mann-Whitney U test). **F,** Frequency (left) and absolute number (right) of pDCs within DCs in the endpoint tdLNs post B16OVA sc challenge (n=10-11 mice per group, from two independent experiments, mean ± SEM, Mann-Whitney U test). (*$p \leq 0.05$, **$p \leq 0.01$, ***$p \leq 0.001$, ****$p \leq 0.0001$)

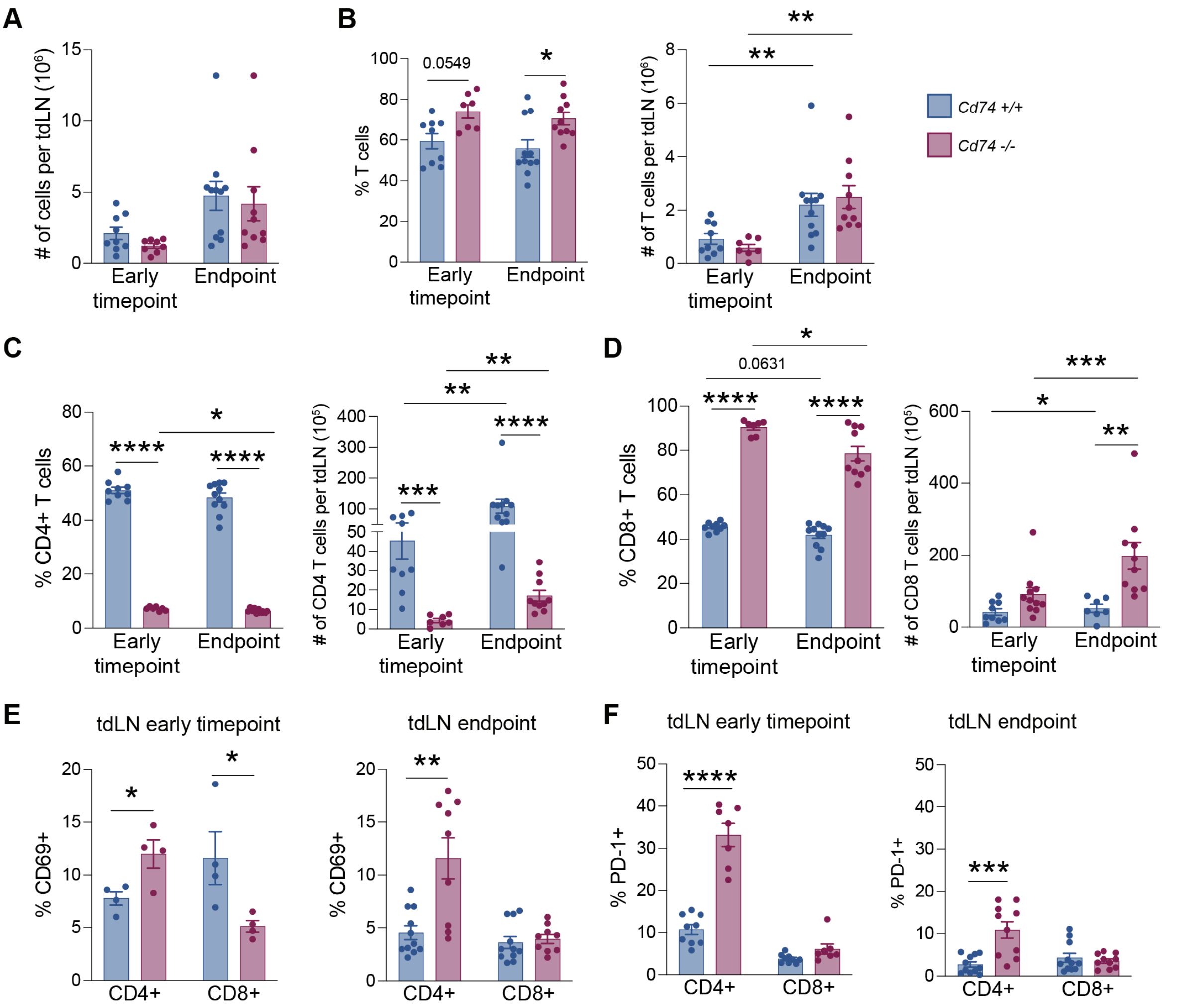


**Fig. S6. Analyses of CD4 and CD8 T cells in the tdLN of *Cd74-/-* and *Cd74+/+* mice in early timepoint and endpoint after B16-OVA challenge. A,** Absolute number of cells per tdLNs at early timepoint and endpoint after tumor challenge (n=8-11 mice per group, from two independent experiments, Mann-Whitney U test). **(B-D)** Frequency (left) and absolute number (right) of B, T cells, C, CD4+, and D, CD8+ T cells in tdLNs in early timepoint and endpoint (n=8-11 mice per group, from two independent experiments, unpaired t-test for frequency results and Mann-Whitney U test for absolute numbers of cells). **E,** Frequency of CD69+ CD4 and CD8 T cells in tdLNs in early timepoint (left) and endpoint (right) (n=8-11 mice per group, from 1-2 independent experiments, unpaired t-test). **F,** Frequency of PD-1+ CD4 and CD8 T cells in tdLNs in early timepoint (left) and endpoint (right) (n=8-11 mice per group, from two independent experiments, unpaired t-test). (*$p \leq 0.05$, **$p \leq 0.01$, ***$p \leq 0.001$, ****$p \leq 0.0001$)

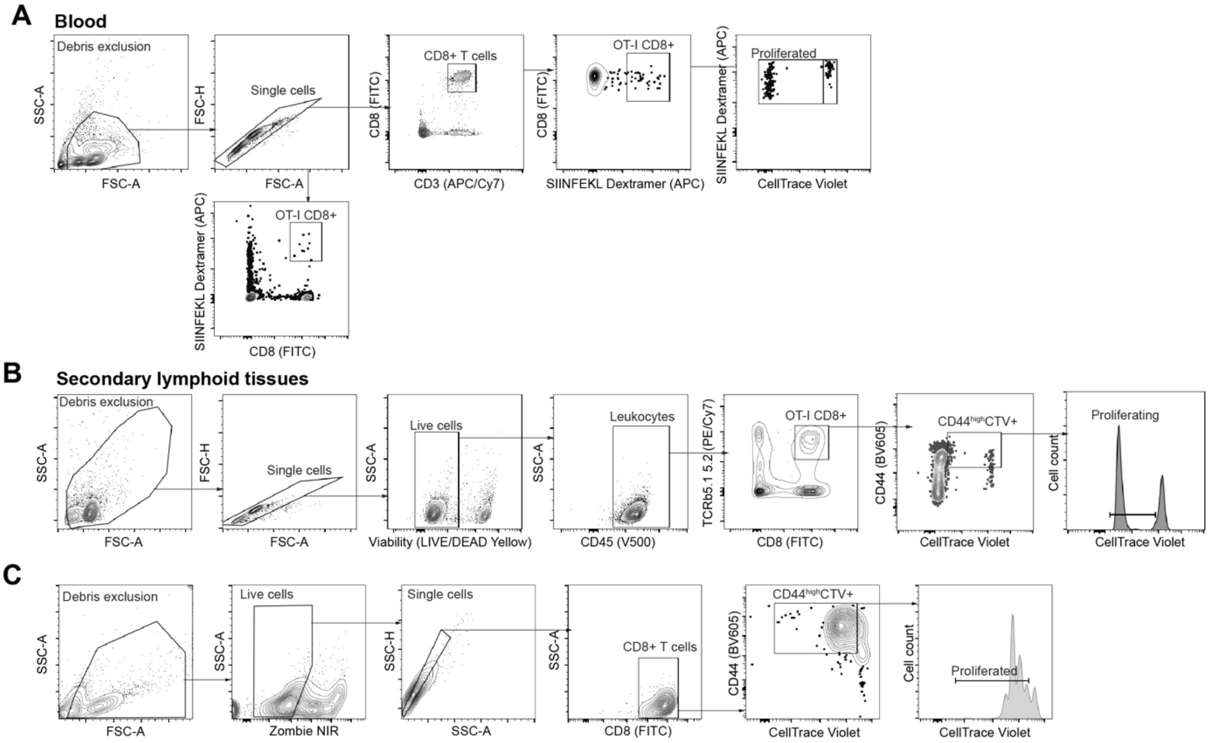


**Fig. S7. Analysis of OT-I CD8+ T cell proliferation. A,** Gating strategy used for the analysis of OT-I CD8+ T cell frequency (within single cells) and proliferation in the periphery of *Cd74 +/+* and *Cd74 -/-* mice. **B,** Gating strategy used for the analysis of OT-I CD8+ T cell frequency and proliferation in the dLN and spleen of *Cd74 +/+* and *Cd74 -/-* mice. **C,** Gating strategy followed for the analysis of OT-I CD8+ T cell proliferation in vitro.

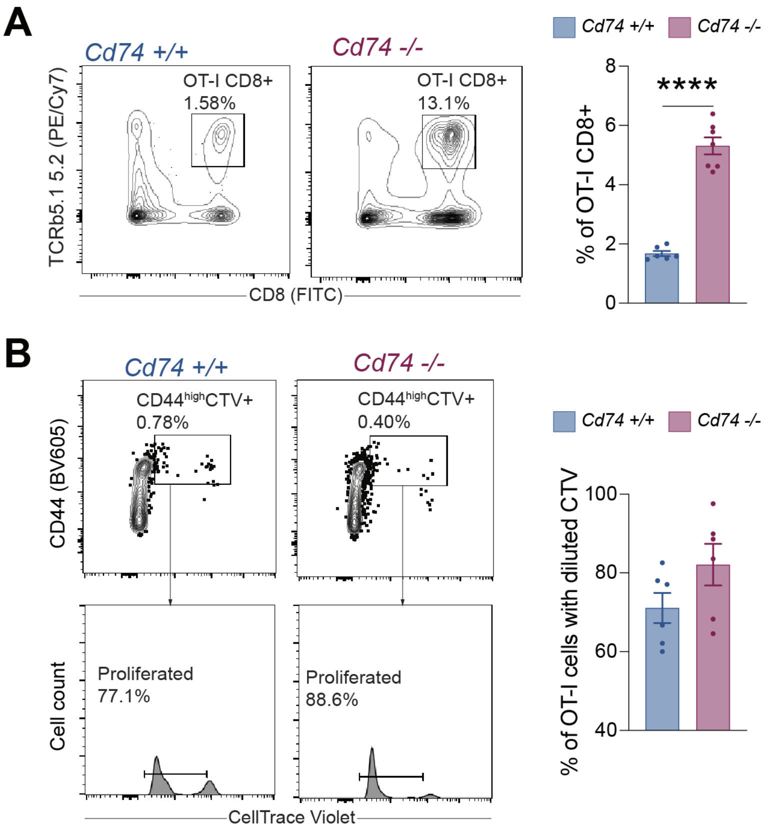


**Fig. S8. Analyses of OT-I CD8+ T cells in spleen of *Cd74 -/-* mice. A,** left: representative flow cytometry contour plots indicating the frequency of OT-I CD8+ T cells in the spleen on day 6 (endpoint), gated within live leukocytes; right: Frequency of OT-I CD8+ T cells in the spleen on day 6 (endpoint) calculated within live leukocytes (n=6 mice per group, from two independent experiments, mean ± SEM, unpaired t-test). **B,** left: Gating strategy used for the discrimination of activated OT-I CD8+ T cells expressing CD44, positive for CTV, and subsequent gating of activated OT-I CD8+ T cells with diluted CTV (proliferating cells); right: Frequency of activated OT-I CD8+ T cells with diluted CTV in the spleen on day 6 (n=6 mice per group, from two independent experiments, mean ± SEM, unpaired t-test). (**p≤0.05*, ***p≤0.01*, ****p≤0.001*, *****p≤0.0001*)

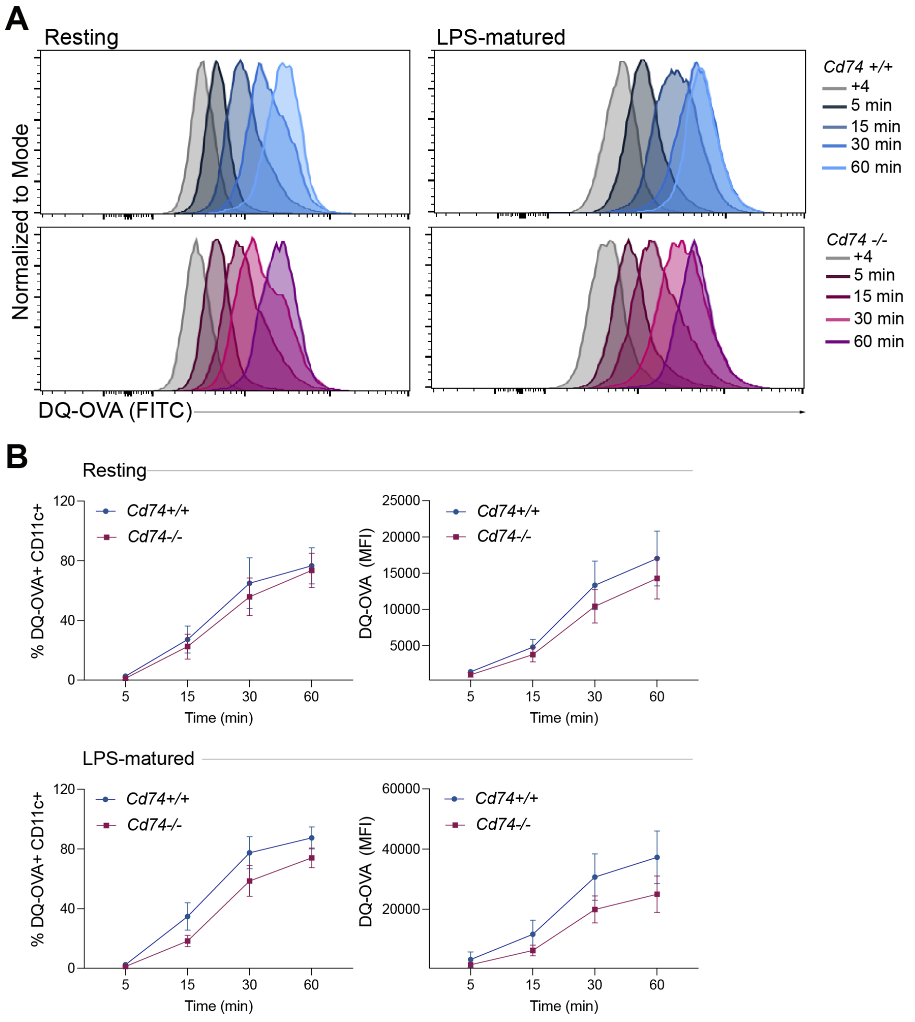


**Fig. S9.** Antigen processing assays with **FL-DCs from *Cd74-/-* mice. A,** Representative flow cytometry histograms of DQ-OVA (40 g/mL) degradation by FL-DCs, resting (left) or LPS-matured (right) for the indicated time points. **B,** Percentage and MFI of DQ-OVA+ (FITC+) FL-DCs in indicated timepoints for resting (top) or LPS-matured (bottom) (n=4 per group, data points indicate FL-DC cultures from individual mice, from 4 independent experiments, mean ± SEM, multiple t-test with Welch's correction).

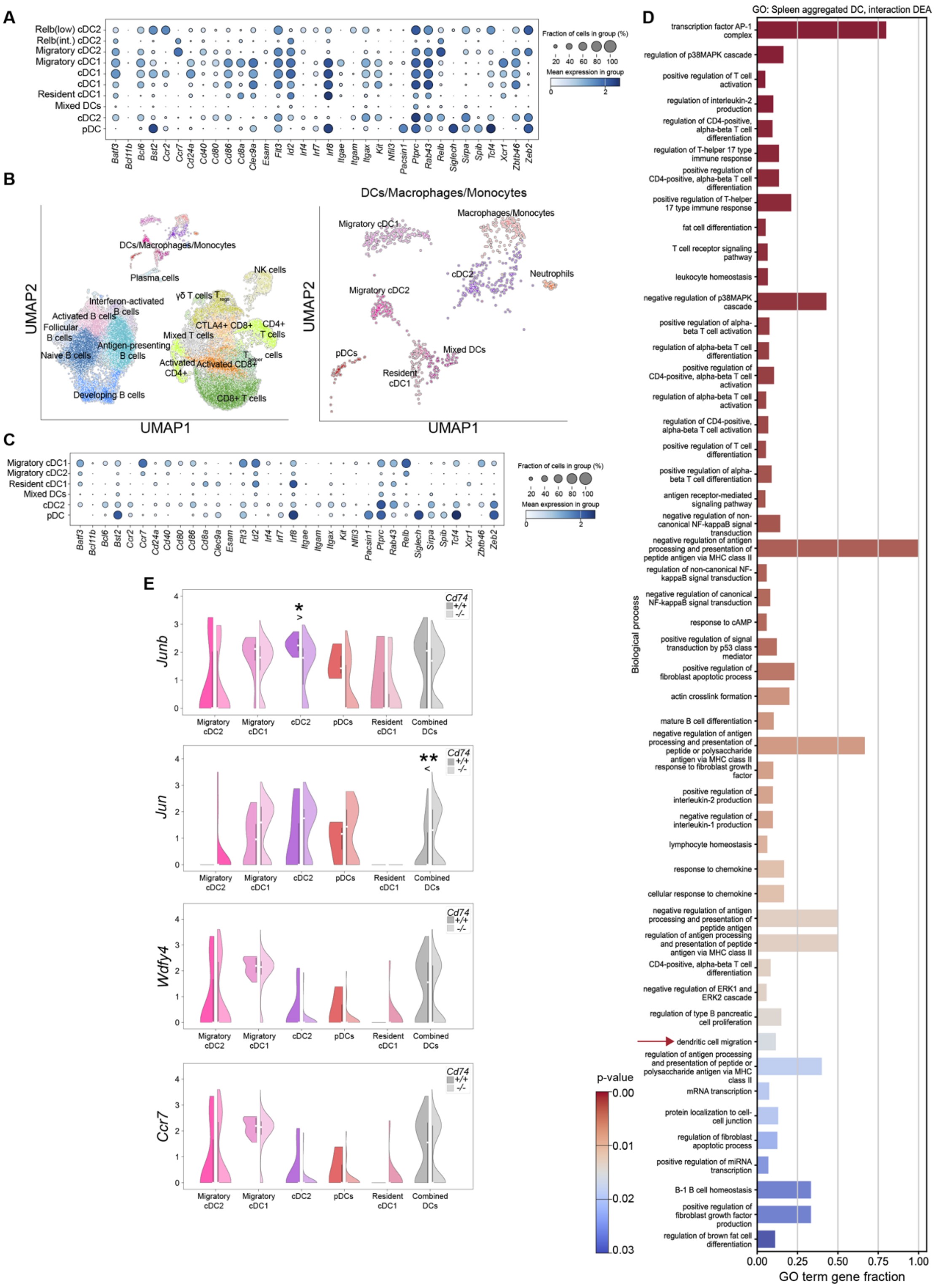


**Fig. S10. Gene markers used to annotate the DC subsets in spleens and LNs and *Jun/Junb* expression in tdLN. A,** Dot plot showing gene markers used to annotate DC clusters in the spleen of naïve and tumor-bearing mice. **B,** UMAPs indicating the cell clusters present in the naïve lymph node and tdLN. **C,** Dot plot showing gene markers used to annotate DC clusters in the lymph node of naïve and tumor-bearing mice. **D,** GO terms with at least 5% of the pathway genes represented in DEGs derived from the interaction effects DEA in the spleen samples of the naïve and B16-OVA-bearing mice (p_corrected<0.03, FDR: Benjamini-Hochberg). **E,** Expression levels (mRNA) of *Junb, Jun*, *Wdfy4*, and *Ccr7* in different DC clusters in the lymph node of tumor-bearing mice (Mann–Whitney U test with Bonferroni correction). (*$p \leq 0.05$, **$p \leq 0.01$, ***$p \leq 0.001$, ****$p \leq 0.0001$)

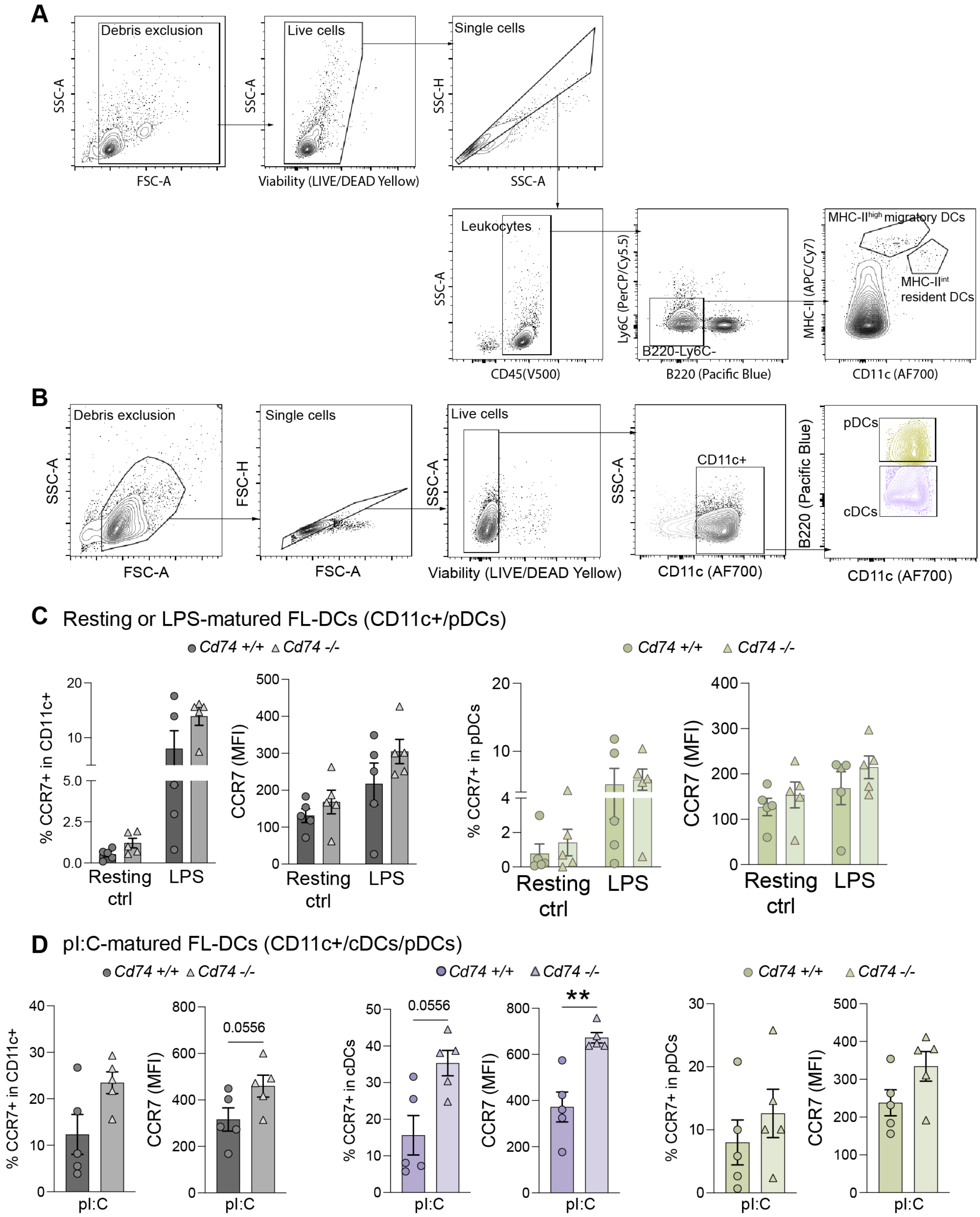


**Fig. S11. Analyses of CCR7 expression on DCs. A,** Gating strategy followed for the evaluation of CCR7 expression on DCs in vivo. **B**, Gating strategy followed to discriminate different populations of DCs generated after differentiation with FLT3L. **C**, CCR7 expression on in vitro generated CD11c+, pDCs (n=5 per group, data points indicate FL-DC cultures from individual mice, from 5 independent experiments, mean ± SEM, Mann-Whitney U test). **D**, CCR7 expression on pI:C-matured in vitro generated CD11c+, cDCs, pDCs (n=5 per group, data points indicate FL-DC cultures from individual mice, from 5 independent experiments, mean ± SEM, Mann-Whitney U test). (**p≤0.05*, ***p≤0.01*, ****p≤0.001*, *****p≤0.0001*)
